# DNA methylation across the genome in aged human skeletal muscle tissue and stem cells: The role of HOX genes and physical activity

**DOI:** 10.1101/2019.12.27.886135

**Authors:** DC Turner, PP Gorski, MF Maasar, RA Seaborne, P Baumert, AD Brown, MO Kitchen, RM Erskine, I Dos-Remedios, S Voisin, N Eynon, RI Sultanov, OV Borisov, AK Larin, EA Semenova, DV Popov, EV Generozov, CE Stewart, B Drust, DJ Owens, II Ahmetov, AP Sharples

**Author notes:** Corresponding author for Skeletal muscle tissue, stem cells and DNA / HOX methylation. Corresponding author for physical activity and HOX methylation. These authors contributed equally to the work (DC Turner, PP Gorski).

## Abstract

Skeletal muscle tissue demonstrates global hypermethylation with aging. However, methylome changes across the time-course of differentiation in aged human muscle derived stem cells, and larger coverage arrays in aged muscle tissue have not been undertaken. Using 850K DNA methylation arrays we compared the methylomes of young (27 ± 4.4 years) and aged (83 ± 4 years) human skeletal muscle and that of young/aged muscle stem cells over several time points of differentiation (0, 72 hours, 7, 10 days). Aged muscle tissue was hypermethylated compared with young tissue, enriched for; ‘pathways-in-cancer’ (including; focal adhesion, MAPK signaling, PI3K-Akt-mTOR signaling, p53 signaling, Jak-STAT signaling, TGF-beta and notch signaling), ‘rap1-signaling’, ‘axon-guidance’ and ‘hippo-signalling’. Aged muscle stem cells also demonstrated a hypermethylated profile in pathways; ‘axon-guidance’, ‘adherens-junction’ and ‘calcium-signaling’, particularly at later timepoints of myotube formation, corresponding with reduced morphological differentiation and reductions in MyoD/Myogenin gene expression compared with young cells. While young cells showed little alteration in DNA methylation during differentiation, aged cells demonstrated extensive and significantly altered DNA methylation, particularly at 7 days of differentiation and most notably in the ‘focal adhesion’ and ‘PI3K-AKT signalling’ pathways. While the methylomes were vastly different between muscle tissue and isolated muscle stem cells, we identified a small number of CpG sites showing a hypermethylated state with age, in both muscle and tissue and stem cells (on genes *KIF15, DYRK2, FHL2, MRPS33, ABCA17P*). Most notably, differential methylation analysis of chromosomal regions identified three locations containing enrichment of 6-8 CpGs in the HOX family of genes altered with age. With *HOXD10, HOXD9, HOXD8, HOXA3, HOXC9, HOXB1, HOXB3, HOXC-AS2* and *HOXC10* all hypermethylated in aged tissue. In aged cells the same HOX genes (and additionally *HOXC-AS3*) displayed the most variable methylation at 7 days of differentiation versus young cells, with *HOXD8, HOXC9, HOXB1* and *HOXC-AS3* hypermethylated and *HOXC10* and *HOXC-AS2* hypomethylated. We also determined that there was an inverse relationship between DNA methylation and gene expression for *HOXB1, HOXA3* and *HOXC-AS3*. Finally, increased physical activity in young adults was associated with oppositely regulating *HOXB1* and *HOXA3* methylation compared with age. Overall, we demonstrate that a considerable number of HOX genes are differentially epigenetically regulated in aged human skeletal muscle and muscle stem cells and increased physical activity may help prevent age-related epigenetic changes in these HOX genes.

## Introduction

Maintaining skeletal muscle mass and function into older age is fundamental for human health-span and quality of life ^1^. Five to ten percent of older humans have sarcopenia ^2, 3^, which is characterized by reductions in muscle mass and strength ^4^. This loss of muscle mass and strength leads to frailty, increased incidence of falls, hospitalization and morbidity ^4, 5, 6, 7, 8, 9, 10, 11^. Annual costs of fragility are estimated to be 39/32 billion (Euros/USD) for European and USA fragility fractures respectively, with the cost of sarcopenia estimated to be £2 billion in the UK ^12^. With an ageing population, these costs are likely to increase with time.

A primary hallmark of ageing is the alteration of the epigenetic landscape. Epigenetics encompasses the interaction between lifestyle/environmental factors and modifications to DNA and histones, without changes to the inherited DNA sequence ^13, 14^. DNA methylation is the most studied epigenetic modification and involves the addition of a covalent methyl group to the 5’ position of the pyrimidine ring of a cytosine (5mC). Increased methylation (hypermethylation) to cytosine-guanine (C-G) pairings (CpG sites), especially in CpG-rich regions such as gene promoters, typically leads to reduced capacity for the transcriptional apparatus to bind to these regions, suppressing gene expression ^14^. Methylated CpG islands in promoters also leads to a tight compaction of adjacent chromatin via the recruitment of chromatin modifying protein/protein-complexes, further silencing gene expression. In contrast, reduced methylation (hypomethylation) provides a more favorable DNA landscape for the transcriptional apparatus to bind to these regions, as well as more ‘relaxed’ chromatin, enabling gene expression to occur.

DNA methylation in aged skeletal muscle occurs at tissue-specific genes ^15^. However, aged muscle also has the smallest overlap with other aged tissue types, suggesting skeletal muscle is unique in comparison with other tissues in its epigenetic aging processes ^15^. Indeed, it has recently been demonstrated that the methylation status of approximately 200 CpG sites can accurately predict chronological age in skeletal muscle tissue ^16^. But that this muscle ‘clock’ only shares 16 of these CpG’s with the original 353 CpG pan-tissue Horvath clock ^16, 17^. Further, using DNA methylation arrays with coverage of ∼450,000 CpG sites ^18^, Zykovich *et al*. demonstrated that compared with young human skeletal muscle, aged skeletal muscle is hypermethylated across the genome. Moreover, our group has demonstrated that mouse skeletal muscle stem cells exposed to a high dose of inflammatory stress in early proliferative life retained hypermethylation of *MyoD* (a muscle-specific regulatory factor) 30 population doublings later ^19^. This suggests that inflamed proliferative aging in muscle stem cells leads to a retained accumulation of DNA methylation. Finally, lifelong physical activity ^20^, endurance and resistance exercise have been associated with predominantly hypomethylation of the genome in young skeletal muscle ^21, 22^. This contrasts with the hypermethylation observed with aging, suggesting that exercise may reverse some age-related changes in DNA methylation.

Skeletal muscle fibers are post-mitotic as they contain terminally differentiated/fused nuclei (myonuclei); thus, repair and regeneration of skeletal muscle tissue is mediated by a separate population of resident stem cells (satellite cells) that can divide. Once activated, satellite cells proliferate and migrate to the site of injury to differentiate and fuse with the existing fibers to enable repair. Target gene analysis showed altered DNA methylation during differentiation of muscle cells into myotubes *in-vitro* ^23^. This included altered methylation of MyoD ^24^, Myogenin ^25^ and Six1 ^26^. While muscle stem cells derived from aged individuals display similar proliferative capacity and time to senescence as young adult cells ^27, 28^, they do have impaired differentiation and fusion into myotubes ^29, 32, 33, 34, 35, 36, 37, 38, 39, 40, 41, 42, 43, 44, 45^. However, a small number of studies did not find an effect of age on the differentiation capacity of isolated cells ^27, 46, 47^. A single study assessed DNA methylation across the genome (450K CpG sites) in aged versus young adult muscle stem cells ^27^ and showed genome-wide hypermethylation in aged cells as well as aged tissue ^27^.

To date, there has been no report of genome-wide DNA methylation dynamics during the entire time-course of muscle cell differentiation, or how age modulates these dynamics. Furthermore, the latest, larger coverage methylation arrays have not yet been implemented in aged muscle tissue. Therefore, the objectives of the current study were: 1) To describe the dynamics of the human DNA methylome in aged and young adult skeletal muscle tissue and primary muscle-derived stem cells over an extensive time-course of differentiation; 0 h (30 minutes post transfer to differentiation media), 72 h (hours), 7 d (days) and 10 d using high coverage 850K CpG arrays. 2) To identify if methylation patterns are similar or different in muscle stem cells compared to skeletal muscle tissue. 3) To test whether increasing physical activity levels is associated with altering DNA methylation in the same genes in aged muscle.

## Methods

### Skeletal muscle biopsies and primary cell isolations

For young adults (n = 9, male, 27 ± 4.4 years-old), skeletal muscle tissue (∼150 mg) was obtained from the vastus lateralis via a conchotome biopsy. Consent and ethical approval were granted for the collection of muscle tissue under NREC, UK approval (16/WM/010) or LJMU, UK local ethics committee approvals [H19/SPS/028 & H15/SPS/031). Six (out of 9) of the young adult’s tissue (male, 28 ± 5.3 years) baseline (at rest) array data was derived from Seaborne *et al*. (2018). This is because we used this baseline tissue to derive cells (detailed below) for DNA methylation analysis of stem cell experiments in the present study. For older adults (n = 5, 2 men/3 women, 83 ± 4 years), tissue biopsies were obtained during elective orthopedic surgeries from University Hospitals of the North Midlands, from the vastus lateralis (knee surgery, n = 2) or gluteus medius muscles (hip surgery, n = 3), under consent and ethical approval 18/WM/0187. DNA and RNA were isolated from these young and aged tissue samples. DNA samples from all 9 young and 5 aged adults were analysed for DNA methylation arrays (detailed below), and a subset were analysed for gene expression (young n = 4, aged n = 5). Primary skeletal muscle cells were derived from a subset of young adult and aged tissue samples, and isolated as per our previous work ^21, 22, 48, 49, 50, 51^. Briefly, approximately 100 mg biopsy tissue was immediately (∼10-30 mins) transferred to a sterile class II biological safety cabinet in pre-cooled (4°C) transfer media (Hams F-10, 2% hi-FBS, 100 U/ml penicillin, 100 μg/ml streptomycin and 2.5 µg/ml amphotericin-B). Any visible connective and adipose tissue were removed using sterile scalpels and muscle tissue was thoroughly washed 2 × in sterile PBS (containing 100 U/ml penicillin, 100 µg/ml streptomycin, 2.5 µg/ml amphotericin-B). PBS was removed and the muscle tissue was minced in the presence of 0.05% Trypsin/0.02% EDTA and all contents (tissue and trypsin) were transferred to a magnetic stirring platform at 37°C for 10 minutes. The addition of 0.05% Trypsin/0.02% EDTA and titration was repeated on any remaining tissue. The supernatant was collected from both procedures and horse serum (HS) was added at 10% of the total supernatant volume to neutralize the trypsin. The supernatant was centrifuged at 340 g for 5 minutes where the supernatant and cell pellet were both plated in separate pre-gelatinised (0.2% gelatin) T25 flasks containing 7.5 ml fresh pre-heated growth media/GM (GM; Ham’s F10 nutrient mix supplemented with 10% hi-NCS, 10% hi-FBS, 100 U/ml penicillin, 100 µg/ml streptomycin, 2.5 µg/ml amphotericin B and 5 mM L-glutamine). Once confluent, cells were trypsinised and reseeded into larger T75’s to expand the cell population. Human derived muscle cells (HMDCs) were seeded onto gelatin coated 6 well plates, at a density of 9 × 10^5^ cells/ml in 2 ml of GM for until ∼90% confluency was attained (∼48 h). GM was removed, and cells underwent 3 × PBS washes before switching to low serum differentiation media (DM; Ham’s F10 nutrient mix supplemented with 2% hiFBS, 100 U/ml penicillin, 100 µg/ml streptomycin, 2.5 µg/ml amphotericin B and 5 mM L-glutamine). HMDCs were differentiated for a total of 10 days (d) which received a 1 ml top up of DM at 72 h and 7 d timepoints. Cells were lysed for DNA and RNA at 0 h (30 minutes in DM), 72 h, 7 d and 10 d, at the same time of day (approx. 08.00-11.00) to minimize the impact of circadian oscillation ^52^. All experiments were carried out below passage 10 to prevent senescence. We undertook methylation arrays on DNA isolated from: 0 h young (n = 7), 0 h aged (n = 4), 72 h young (n = 7), 72 h aged (n = 4), 7 d young (n= 6), 7d aged (n = 3), 10 d young (n = 2) and 10 d aged (n= 4). Gene expression was analysed using young (n= 4, 7 d) and aged (n = 3, 7 d) cells. It’s worth noting that we had n= 3-4 for most conditions, however in the 10 d young cells condition, the DNA did not pass QA/QC for the arrays, and we had no cells left for these participants. Therefore, unfortunately we could only run n = 2 for this single condition. Therefore, results for this condition should be viewed with this caveat in mind. A schematic of experimental design can be found in Suppl. Figure **1**.

### Myogenicity and morphology measurements

Following isolations, attached single cells were fixed prior to staining and fluorescent immunocytochemistry analysis to determine myogenicity (via desmin positivity) of the isolated young and aged muscle derived cultures. Briefly, approximately 2 × 10^4^ cells were seeded onto 3 × wells of a 6-well plate and were incubated for 24 h in GM. Existing media was removed and cells were washed 3 × in PBS before fixation using the graded methanol/acetone method (50:25:25 TBS:methanol:acetone for 15 minutes followed by 50:50 methanol:acetone) after which cells were permeabilised in 0.2% Triton X-100 and blocked in 5% goat serum (Sigma-Aldrich, UK) in TBS for 30 minutes. Cells were washed 3 × in TBS and incubated overnight (4 °C) in 300 μl of TBS (with 2% goat serum and 0.2% Triton X-100) containing primary anti-desmin antibody (1:50; ab15200, Abcam, UK). After overnight incubation, cells were washed 3 × in TBS and incubated at RT for 3 h in 300 μl of secondary antibody solution (TBS, 2% goat serum and 0.2% Triton X-100) containing the secondary antibody, anti-rabbit TRITC (1:75; T6778, Sigma-Aldrich, UK) to counterstain desmin. Finally, cells were washed again 3 × in TBS, prior to counterstaining nuclei using 300 μl of DAPI solution at a concentration of (300 nM; D1306, Thermo Fisher Scientific, UK) for 30 minutes. Immunostained cells were then visualised using a fluorescent microscope (Nikon, Eclipse Ti-S, Japan or Olympus IX83, Japan) and imaged using corresponding software (Nikon, NIS Elements and Olympus FV10-ASW 4.2). Myoblasts, myotubes and nuclei were visualized using TRITC (Desmin, Excitation: 557 nm, Emission: 576 nm) DAPI (Excitation: 358 nm, Emission: 461 nm) filter cubes. All immunostained samples were imported to Fiji/ImageJ (version 2.0.0) software for subsequent calculations. Note, there was no difference (p = 0.86) in myogenic cell proportions in aged (35 ± 5%) versus young cells (34 ± 9%). Therefore, the aged and young muscle derived cells were matched for their desmin positivity (% myoblasts) prior to differentiation experiments and downstream DNA methylation and gene expression analysis. To determine myotube differentiation and formation at each time point, cells were imaged using light microscopy (Olympus, CKX31, Japan) at 0, 72 h, 7 and 10 d of differentiation.

### DNA isolation and bisulfite conversion

Prior to DNA isolation, tissue samples were homogenized for 45 seconds at 6,000 rpm × 3 (5 minutes on ice in between intervals) in lysis buffer (180 µl buffer ATL with 20 µl proteinase K) provided in the DNeasy spin column kit (Qiagen, UK) using a Roche Magnalyser instrument and homogenization tubes containing ceramic beads (Roche, UK). DNA was then isolated using the DNAeasy kit (Qiagen, UK) according to manufacturer’s instructions. Cells were lysed in 180 µl PBS containing 20 µl proteinase K, scraped from wells, and then isolated using DNAeasy kits (Qiagen, UK) as above with the tissue lysates. The DNA was then bisulfite converted using the EZ DNA Methylation Kit (Zymo Research, CA, United States) as per manufacturer’s instructions.

### Infinium MethylationEPIC BeadChip Array

All DNA methylation experiments were performed in accordance with Illumina manufacturer instructions for the Infinium MethylationEPIC 850K BeadChip Array (Illumina, USA). Methods for the amplification, fragmentation, precipitation and resuspension of amplified DNA, hybridisation to EPIC beadchip, extension and staining of the bisulfite converted DNA (BCD) can be found in detail in our open access methods paper ^49^. EPIC BeadChips were imaged using the Illumina iScan® System (Illumina, United States).

### DNA methylation analysis, CpG enrichment analysis (GO and KEGG pathways), differentially modified region analysis and Self Organising Map (SOM) profiling

Following DNA methylation quantification via MethylationEPIC BeadChip array, raw .IDAT files were processed using Partek Genomics Suite V.7 (Partek Inc. Missouri, USA) and annotated using the MethylationEPIC_v-1-0_B4 manifest file. We first checked the average detection p-values. The highest average detection p-value for the samples was 0.0023 (Suppl. Figure **2a**), which is well below the recommended 0.01 in the Oshlack workflow ^53^. We also produced density plots of the raw intensities/signals of the probes (Suppl. Figure **2b**). These demonstrated that all methylated and unmethylated signals were over 11.5 (mean was 11.52 and median was 11.8), and the difference between median methylation and median unmethylated signal was 0.56 ^53^. Upon import of the data into Partek Genomics Suite we removed probes that spanned X and Y chromosomes from the analysis due to having both males and females in the study design, and although the average detection p-value for each samples was on average very low (no higher than 0.0023) we also excluded any individual probes with a detection p-value that was above 0.01 as recommended in ^53^. Out of a total of 865,860 probes, removal of the X and Y chromosome probes and those with a detection p-value above 0.01 reduced the probe number to 846,233 (removed 19,627 probes). We also filtered out probes located in known single-nucleotide polymorphisms (SNPs) and any known cross-reactive probes using previously defined SNP and cross-reactive probe lists identified in earlier 850K validation studies ^54^. This resulted in a final list of 793,200 probes to be analysed. Following this, background normalisation was performed via functional normalisation (with noob background correction), as previously described ^55^. Following functional normalisation, we also undertook quality control procedures of principle component analysis (PCA), density plots by lines as well as box and whisker plots of the normalised data for tissue samples (Suppl. Figures **2c, d, e** respectively) and cell samples (Suppl. Figures **2 f, g, h** respectively). Any outlier samples were detected using PCA and the normal distribution of β-values. Outliers were then removed if they fell outside 2 standard deviations (SDs) (e.g. Suppl. Figure **2c**) of the ellipsoids and/or if they demonstrated different distribution patterns to the samples of the same condition. Only one young adult male tissue sample was removed due to being outside 2 standard deviations outside samples from the same condition (Suppl. Figure **2c**; sample with a strikethrough line). Following normalisation and quality control procedures, we undertook differential methylation analysis by converting β-values to M-values (M-value = log2(β / (1 - β)), as M-values show distributions that are more statistically valid for the differential analysis of methylation levels ^56^. We undertook a 1-way ANOVA for comparisons of young and aged skeletal muscle tissue. For primary human muscle cells, we undertook a 2-way ANOVA for age (young vs. aged cells) x time (0, 72 h, 7 and 10 d) and also explored the ANOVA main effects for both age and time independently. We also performed planned contrasts within the differentiation time-course in both young adult cells alone and aged cells alone e.g. 0 h vs. 72 h, 0 h vs. 7 d 0 h vs. 10 d, as well as planned contrasts for young vs aged cells at each time point of differentiation (0 h vs. 0 h, 72 h vs. 72 h, 7 d vs. 7 d, 10 d vs. 10 d in aged vs. young adult cells respectively). Any CpG with a False Discovery Rate (FDR) ≤ 0.05 was deemed significant. In some analyses, e.g. > 100,000 differentially methylated CpG sites were identified at FDR ≤ 0.05, so we also show results at a more stringent FDR of ≤ 0.01 or 0.001, or at FDR of ≤ 0.05 and ‘a change (difference in M-value) in methylation greater than 2’. These shorter lists of CpGs contain the most significant sites to enable sensible pathway analysis. We specify when a more stringent FDR than 0.05 was used in the results text. We then undertook CpG enrichment analysis on these differentially methylated CpG lists within gene ontology and KEGG pathways ^57, 58, 59^ using Partek Genomics Suite and Partek Pathway. We also undertook differentially methylated region analysis (e.g. identifies where several CpGs are consistently differentially methylated within a short chromosomal location/region) using the Bioconductor package DMRcate (DOI: 10.18129/B9.bioc.DMRcate). Finally, in order to plot temporal changes in methylation across the time-course of muscle cell differentiation we undertook Self Organising Map (SOM) profiling of the change in mean methylation within each condition using Partek Genomics Suite.

### RNA isolation, primer design & gene expression analysis

Skeletal muscle tissue muscle was homogenised in tubes containing ceramic beads (MagNA Lyser Green Beads, Roche, Germany) and 1 ml Tri-Reagent (Invitrogen, Loughborough, UK) for 45 seconds at 6,000 rpm × 3 (and placed on ice for 5 minutes at the end of each 45 second homogenization) using a Roche Magnalyser instrument (Roche, Germany). Human cells from differentiation experiments were also lysed in 300 µl Tri-Reagent for 5 minutes at RT mechanically dissociated/lysed using a sterile scraper. RNA was then isolated as per Invitrogen’s manufacturer’s instructions for Tri-reagent. Then a one-step RT-PCR reaction (reverse transcription and PCR) was performed using QuantiFastTM SYBR Green RT-PCR one-step kits on a Rotorgene 3000Q. Each reaction was setup as follows; 4.75 μl experimental sample (7.36 ng/μl totalling 35 ng per reaction), 0.075 μl of both forward and reverse primer of the gene of interest (100 μM stock suspension), 0.1 μl of QuantiFast RT Mix (Qiagen, Manchester, UK) and 5 μl of QuantiFast SYBR Green RT-PCR Master Mix (Qiagen, Manchester, UK). Reverse transcription was initiated with a hold at 50°C for 10 minutes (cDNA synthesis) and a 5-minute hold at 95°C (transcriptase inactivation and initial denaturation), before 40-50 PCR cycles of; 95°C for 10 sec (denaturation) followed by 60°C for 30 sec (annealing and extension). Primer sequences for genes of interest and reference genes are included in Table 1. All genes demonstrated no unintended targets via BLAST search and yielded a single peak after melt curve analysis conducted after the PCR step above. All relative gene expression was quantified using the comparative Ct (^ΔΔ^Ct) method ^60^. For human cell differentiation analysis via measurement of myoD and myogenin, a pooled mean Ct for the 0 h young adult control samples were used as the calibrator when comparing aged vs. young adult cells. This approach demonstrated a low % variation in Ct value of 9.5 and 8.5% for myoD and myogenin, respectively. For HOX gene analysis between young and aged tissue and for the 7 d aged cells vs. 7 d young adult cells, the mean Ct of the young adult cells were used as the calibrator. The average, standard deviation and variations in Ct value for the B2M reference gene demonstrated low variation across all samples (mean ± SD, 13.12 ± 0.98, 7.45% variation) for the analysis of myoD and myogenin. For HOX gene analysis, the RPL13a reference gene variation was low in the human tissue (17.77 ± 1.71, 9.6% variation) and stem cell (15.51 ± 0.59, 3.82% variation) experiments. The average PCR efficiencies of myoD and myogenin were comparable (94.69 ± 8.9%, 9.4% variation) with the reference gene B2M (89.45 ± 3.76%, 4.2% variation). The average PCR efficiencies of the all genes of interest in the tissue analysis of the HOX genes were also comparable (90.87 ± 3.17%, 3.39% variation) with the human reference gene RPL13a (92 ± 2.67%, 2.9% variation). Similarly, for the cell analysis, HOX genes of interest efficiencies were comparable (89.59 ± 4.41%, 4.93% variation 4.93%) with the reference gene RPL13a (89.57 ± 3.55%, 3.97% variation). Statistical analysis for HOX genes was performed using t-tests (aged tissue vs. young tissue and 7d aged versus 7d young).

### Physical Activity and DNA methylation

The human association study involved 30 physically active and endurance-oriented men of Eastern European descent (32.9 ± 9.9 years). The study was approved by the Ethics Committee of the Physiological Section of the Russian National Committee for Biological Ethics and Ethics Committee of the Federal Research and Clinical Center of Physical-chemical Medicine of the Federal Medical and Biological Agency of Russia. Written informed consent was obtained from each participant. The study complied with the guidelines set out in the Declaration of Helsinki. Physical activity was assessed using questionnaire. Participants were classified as mildly active (1-2 training sessions per week, n=6), moderately active (3-4 training sessions per week, n=8) and highly active (5-7 sessions per week, n=16) participating in aerobic exercise for at least the last 6 months. Bisulfite conversion of genomic DNA was performed using the EpiMark® Bisulfite Conversion Kit in accordance with the manufacturer’s instructions. In the same analysis as described above for the aged muscle tissue and stem cell data, methylome of the vastus lateralis in the physically active men was evaluated using the Infinium MethylationEPIC 850K BeadChip Array (Illumina, USA) and imaged and scanned using the Illumina iScan® System (Illumina, United States). Also as in the above analysis we filtered out probes located in known single-nucleotide polymorphisms (SNPs) and any known cross-reactive probes using previously defined SNP and cross-reactive probe lists ^61^. Low quality probes were also filtered, as with the above analysis. The final analyses included 796,180 of 868,565 probes. Data were normalised using the same functional normalisation (with noob background correction), as in the aged tissue and stem cell data, and as previously described ^55^. Any outliers were interrogated via PCA as above, however all samples passed the QC and therefore there were no outliers. The methylation level of each CpG-site after normalization and filtering processes was represented as a β-value ranging from 0 (unmethylated) to 1 (fully methylated) in order to undertake multiple regression (as regression analysis performed better with finite values. Ideally, if values range from 0 to 1 with beta-values satisfying this criteria). Statistical analyses were conducted using PLINK v1.90, R (version 3.4.3) and GraphPad InStat (GraphPad Software, Inc., USA) software. Multiple regression was used for testing associations between the CpG-methylation level and physical activity adjusted for age and muscle fibre composition. With methylation level as the dependent variable, and physical activity and age as independent variables. P values < 0.05 were considered statistically significant to test the identified HOX CpG sites. The p value is given for the physical activity score adjusted for age and muscle fibre composition.

## Results

### Aged human skeletal muscle tissue is hypermethylated compared with young adult tissue

There were 17,994 differentially methylated CpG positions (DMPs) between aged and young adult tissue at FDR ≤ 0.05 (Suppl. File **1a**), and 6,828 DMPs at a more stringent FDR of ≤ 0.01 (Suppl. File **1b**). The overwhelming majority of the 6,828 DMPs (93%) were hypermethylated in aged compared with young muscle (Figure **1a**). Furthermore, DMPs were enriched in CpG islands (5,018 out of 6,828 CpG’s) with the remaining in N-shores (354), S-Shelf (48), S-Shores (341), S-Shelf (69) and ‘other’ (998). Ninety nine percent of the DMPs (4,976 out of the 5,018 CpGs) located in CpG islands were also hypermethylated with age compared to young adult tissue (Suppl. File **1c**). After gene ontology analysis on these DMPs (Suppl. File **1d)**, hypermethylation was enriched within the three overarching gene ontology terms: ‘Biological process’ (Suppl. Figure **3a**), ‘cellular component;’ (Suppl. Figure **3b**), and ‘molecular function’ (Suppl. Figure **3c**). Within the GO terms that included the search term ‘muscle’, the most significantly enriched was ‘regulation of muscle system process’ (Figure **1b**, CpG list Suppl. File **1e**). Within this GO term was ‘regulation of muscle contraction’ (CpG list Suppl. file **1f**). Further, we found hypermethylation enrichment in KEGG pathways: ‘Pathways in cancer’, ‘Rap1 signaling’, ‘Axon guidance’ and ‘Hippo signaling’ (Suppl. File **1g**). Within the top enriched pathway, ‘pathways in cancer’, 96% (266 out of 277 CpGs) were hypermethylated and only the 4% (11 CpGs) were hypomethylated in aged compared with young adult muscle tissue (Suppl. Figure **4**; Suppl. File **1h**). Differentially methylated regions (DMRs) were also analysed between young and aged skeletal muscle tissue (Suppl. File **1i**). The top 5 DMRs were identified as: chr8:22422678-22423092 (415 bp) within a CpG island of the SORBS3 gene with 6 CpG’s that were all hypermethylated in aged versus young adult tissue. Also, chr6:30653732-30655720 (1989 bp) within a CpG island of the PPPR1 gene contained 8 CpG’s that were all hypermethylated compared with young adult muscle. Chr20:13200939-13202437 (1499 bp) within a CpG island of the promoter of gene ISM, contained 8 CpG sites were also all hypermethylated. Similarly, the gene PDE4D1P on Chr1:144931376-144932480 (1105 bp) contained 6 CpG sites that spanned its promoter within a CpG island, once again demonstrating hypermethylation. The only gene demonstrating the opposite direction in methylation status (hypomethylation) in aged tissue was the gene C1orf132 (new gene name MIR29B2CHG), also on chromosome 1 (Chr1:207990896-207991936; 1041 bp), where 5 CpGs within this region were hypomethylated in opposition to the majority of gene regions that were hypermethylated in aged versus young tissue.

**Figure 1.**
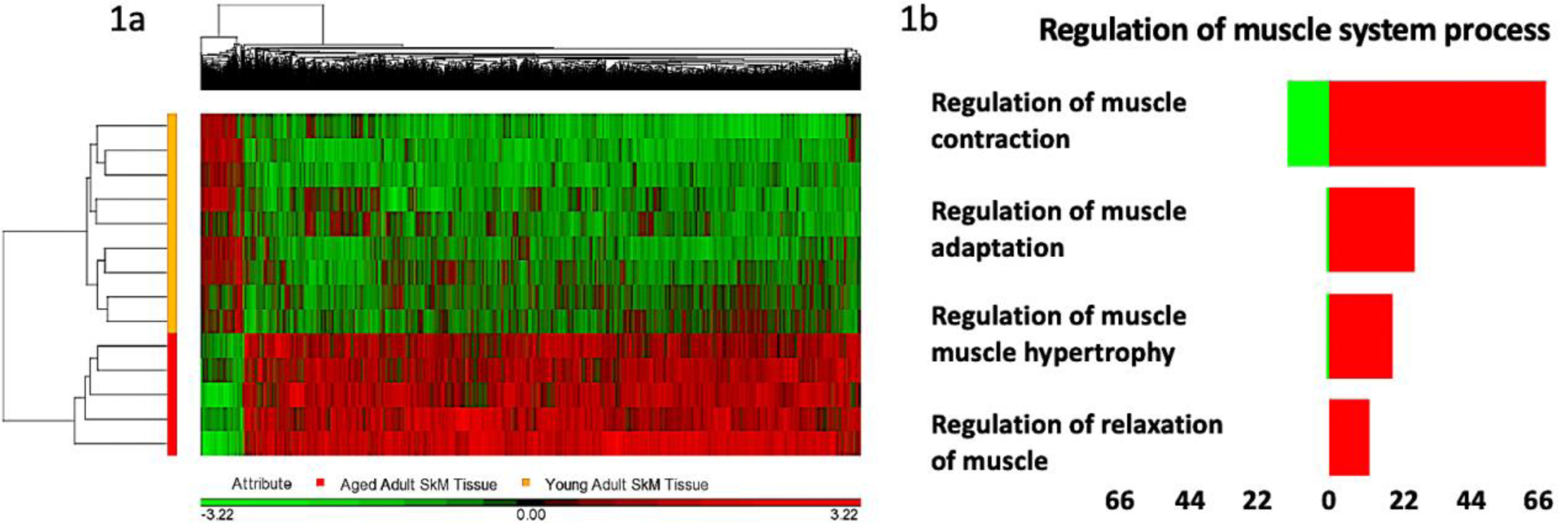
DNA methylation across the genome in aged human skeletal muscle tissue compared with young adult tissue. **1a**. Hierarchical clustering heatmap of the significantly differentially methylated CpG sites, depicting aged skeletal muscle containing a hypermethylated (RED) versus hypomethylated (GREEN) signature compared with young adult tissue. **1b**. Significantly enriched CpG sites for most significantly enriched GO term containing the search term ‘muscle’; ‘regulation of muscle system process’ (RED hypermethylated and GREEN hypomethylated CpGs).

### Aged primary human muscle stem cells displayed more varied DNA methylation signatures versus young adults during the time-course of differentiation

We analysed DNA methylation from differentiating human muscle stem cells at 0 h (30 minutes post dropping to 2% serum), 72 h, 7 and 10 d. Both young and aged adult cells demonstrated morphological differentiation and myotube formation as the cells advanced across the time-course (Figure **2a**). This was confirmed by increases in myoD and myogenin mRNA expression in both young and aged cells as differentiation progressed up to 72hrs (Figure **2b**). However, myotube formation was less extensive in elderly cells, despite young and aged cells having identical numbers of starting myogenic cells. This age associated reduction in differentiation was confirmed with significantly reduced myoD and myogenin gene expression at 72 h and 10 d (p ≤ 0.05), as well as by delayed increases in myogenin mRNA expression (Figure **2b**) in aged cells. A similar delay has been shown previously to be associated with impairing the fusion process and myotube formation ^32, 33^. There were also differences in DNA methylation between aged and young cells during differentiation. Indeed, the interaction for a 2-way ANOVA (Age × Time) generated a list of 40,854 CpG sites that were significantly altered with age and across the time-course of differentiation (FDR ≤ 0.05, Suppl. File **2a**). With a more stringent FDR ≤ 0.01 and ≤ 0.001 there were still 9,938 and 2,020 CpG’s significantly differentially methylated respectively in aged cells versus young cells across all time points of differentiation (Suppl. File **2b** and **2c** respectively).

**Figure 2.**
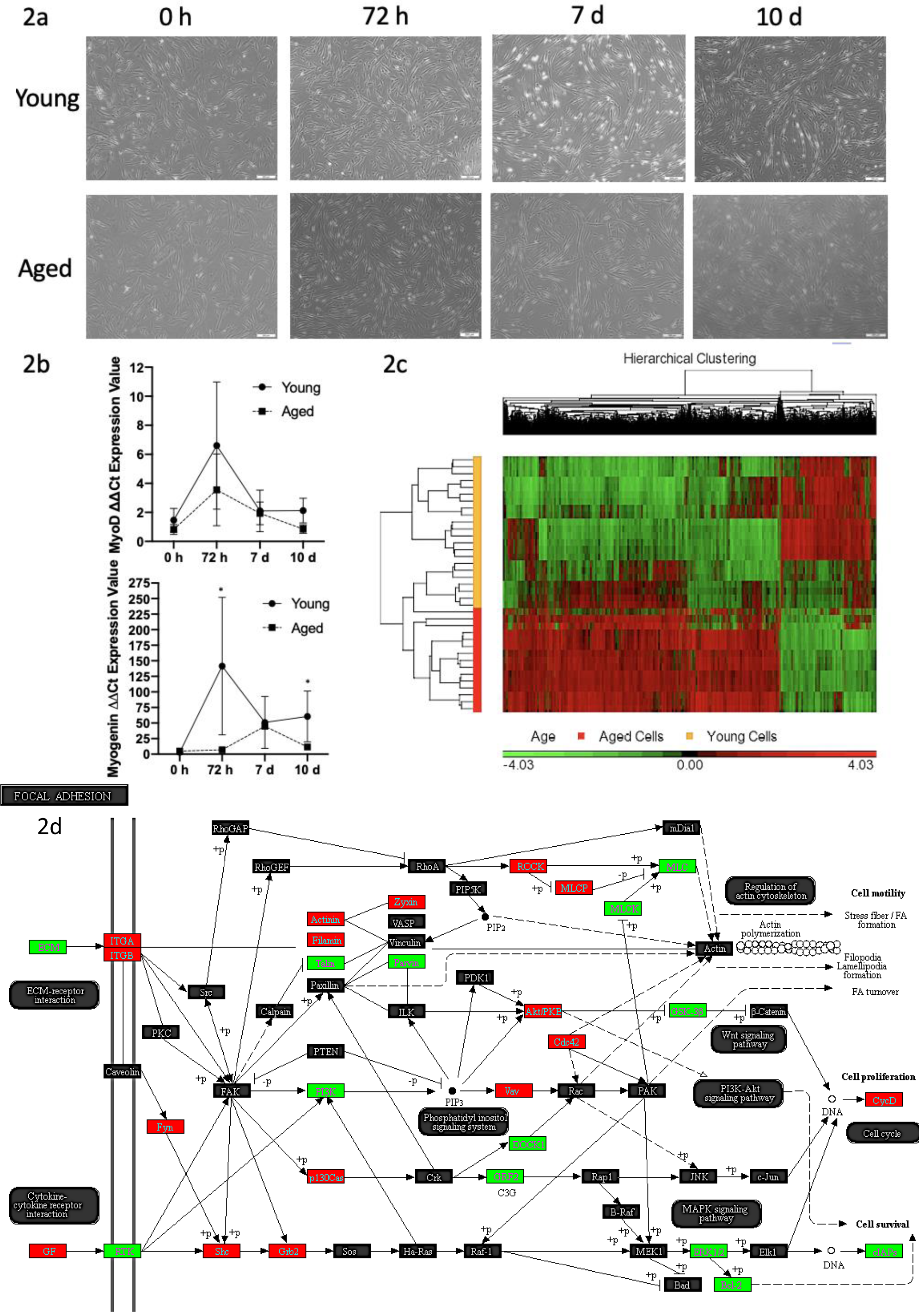
Aged and young adult primary muscle derived stem cells differentiated over 0, 72 h, 7d and 10 d and differences in DNA methylation. **a**. Light microscope images of aged versus young muscle derived stem cells, depict fewer myotubes in aged versus young cells, particularly at 7 and 10 days. **b**. MyoD and myogenin gene expression in aged versus young cells over the differentiation time course. Where, reductions in myogenin were observed at 72 h in aged cells as well as a delayed increase in the upregulation of myogenin compared with young cells. **c**. Hierarchical clustering heatmap of the significantly differentially methylated CpG sites between aged and young muscle stem cells across the entire time-course of differentiation (all time points of 0, 72 h, 7 d and 10 d). RED hypermethylated and GREEN hypomethylated. **d**. Comparison of DNA methylation in aged versus young muscle stem cells at 7 d of differentiation (7 d aged vs. 7 d young adult cells) in most enriched KEGG pathway, ‘focal adhesion’. Note, because some genes have multiple CpGs per gene, Partek Pathway (Partek Genomics Suite) used to create figure 2d, selects the CpG with the lowest p-value (most significant) and uses the beta-value for that particular CpG to colour the pathway diagram. This is therefore accurate in situations where CpGs on the same gene have the same methylation pattern (e.g. all hyper or hypomethylated). However, where multiple CpGs on the same gene have the different methylation pattern (e.g. some hypomethylated and others hypermethylated), only the most significant methylated CpG is chosen and represented in the image. For full and accurate significant CpG lists, including the sites that were hypo and hypermethylated in this ‘focal adhesion’ pathway, are included in Suppl. File 5i.

We next assessed changes in DNA methylation across time during differentiation (main effect for ‘time’) i.e. changes in DNA methylation overtime time that was visible in both young and old cells). However, we did not find time-related DMPs at our statistical cut off of FDR ≤ 0.05. This suggested, that DNA methylation was not considerably changing over the time course of muscle stem cell differentiation itself. We therefore contrasted each timepoint of differentiation with the baseline (0 h timepoint) in young and aged cells to further examine this observation. Indeed, in young adult cells, there were only 1 and 14 DMPs at 72 h and 10 d respectively (FDR ≤ 0.05), and no DMPs at 7 d (FDR ≤ 0.05) compared to their baseline 0 h timepoint. In aged cells, there were no DMPs during early differentiation (72 h vs. 0 h, FDR ≤ 0.05) but 2,785 significant DMPs at 7 days of differentiation (7d vs. 0 h, FDR ≤ 0.05, Suppl. file **2d**) and 404 DMPs at 10 days of differentiation (10 d vs. 0 h, FDR ≤ 0.05, Suppl. File **2e**). Therefore, while the differentiation itself was not changing DMPs in young cells there were a significant number of DMPs at 7 days of differentiation in aged cells compared with their own 0 h timepoint. Using more stringent cut-offs (FDR ≤ 0.05 and change/difference in methylation greater than 2), at this 7-day time point in aged cells there were 1,229 DMPs at 7 d vs 0 h (Suppl. File **2f**), with a balanced number of hypo and hypermethylated DMPs. Overall, this may suggest a more dysfunctional DNA methylation program during differentiation in aged human muscle cells compared with young cells. Conducting gene ontology analyses at this 7 day time point (Suppl. File **2g**), it suggested that this varied methylation response in aged cells was enriched in GO terms: ‘regulation of localisation’, ‘regulation of cell communication’ and ‘regulation of signaling’ (Suppl. Figure **5**; Suppl. File **2h, i, j** respectively). CpGs within these ontologies also confirmed that there was a similar hypo and hypermethylation profile in aged cells at 7 days. Further, KEGG pathway analysis suggested the top enriched pathways at 7 days in aged cells versus 0 h were: ‘axon guidance’, cholinergic synapse’, ‘adrenergic signaling in cardiomyocytes’ and ‘circadian entrainment’ (Suppl. File **2k**). Finally, DMR analysis between young and aged cells at 7 days of differentiation identified two regions in 2 genes that had 5 or more CpG sites significantly differentially methylated (Suppl. File **2l**). The first being Chr3:155394096-155394303 (209 bp) located on gene PLCH1 that had 5 of its CpG’s hypomethylated. The second being a region non-annotated in location Chr2: 119615888-119617128 (1241 bp) that was hypermethylated on 5 CpG’s. Suggesting that enriched and varied methylation of these regions occurred at 7 d differentiation in aged cells that was not detected at 7 days in young adult cells.

### Aged muscle cells demonstrate hypermethylated signatures versus young muscle cells across differentiation, particularly at 7 days

We next wished to identify differences in DNA methylation in aged versus young muscle stem cells. The main effect for ‘age’ generated a significant differentially methylated (FDR ≤ 0.05) CpG list of 269,898 sites that significantly varied in aged cells versus young cells. Even with a more stringent cut-off (FDR ≤ 0.01) there were still 159,235 sites significantly modified (Suppl. File **3a**). Increasing the stringency to a difference of greater than 2 while keeping an FDR ≤ 0.01 identified 2,719 DMPs with the most highly significant and demonstrated the largest differences between aged and young cells (Suppl. File **3b**). As with the aged versus young skeletal muscle tissue analysis, the majority of these significantly modified CpG sites in aged cells were hypermethylated (2,017 out of 2,719) versus hypomethylated (702 out of 2,719) compared with young cells, visualised in hierarchical clustering aged cells vs. young cells (Figure **2c**). Two hundred and eleven out of these 2,719 CpGs were located in islands and 97 were promoter associated. Gene ontology identified that aged cells demonstrated this significantly enriched hypermethylated profile in GO terms: ‘developmental process’, ‘anatomical structure development’, ‘anatomical structure morphogenesis’ (Suppl. File **3c)** and also in KEGG pathways ‘axon guidance’, adherens junction’, ‘calcium signaling’, ‘focal adhesion’ and ‘protein digestion and absorption’ (Suppl. File **3d)**. DMR analysis (Suppl. File **3e**) identified that a non-coding location on chr12:115134344-115135576 (1233 bp) that contained 13 CpG’s that were hypermethylated in aged cells versus young cells. Further, there were 7 CpG’s that were hypermethylated in the region of the gene LY6G5C (Chr6:31650736-31651158, 423 bp). There was also region containing 8 CpGs of the HOXC10 gene just upstream of HOXC6 and MIR196 (Chr12:54383692-54385621, 1933 bp) that all demonstrated a hypomethylated profile in aged cells vs. young cells. Interestingly, on the same chromosome just upstream of the HOXC10 gene (Chr12:54376020-54377868, 1849 bp), within the lncRNA HOXC-AS3, there were another 6 CpGs that were hypomethylated in aged cells versus young cells. The concentration of hypomethylated CpGs in aged versus young cells in HOXC genes is interesting given the majority of CpG’s, more generally, were hypermethylated in aged cells versus young cells. A finding that is taken further in the later results section below.

Given these changes, predominantly favouring hypermethylation (except the HOXC genes identified above) in aged cells versus young cells, we also performed contrasts between aged and young adult cells at each time point of differentiation. When we compared 0 h aged cells with young adult cells at 0 h, we identified 738 DMPs between young and old cells (FDR ≤ 0.05, with a difference/change greater than 2), 79% of which were hypermethylated in aged versus young cells (Suppl. File **3f**). Gene ontology analysis (Suppl. File **3g)**revealed that the DMPs were in genes enriched for ‘cytoskeletal protein binding’ (Suppl. Figure **6a;** Suppl. File **3h**), ‘developmental process’, ‘cell junction’, ‘cytoskeleton’ and ‘actin binding’ (Suppl. File **3i, j, k, l** respectively). KEGG pathway analysis of 0 h aged versus 0 hr young adult cells (Suppl. File **3m**) also suggested that there were significantly enriched hypermethylated pathways for ‘Axon Guidance’ (Suppl. Figure **6b**; CpG List Suppl. File **3n**), ‘Insulin secretion’, ‘Phospholipase D signaling’, ‘cAMP signaling pathway’ and ‘Aldosterone synthesis and secretion’ (Suppl. Files **3o, p, q, r** respectively). At 72 h d, there were 1,418 DMPs between aged and young adult cells (FDR ≤ 0.05), and with a difference/change greater than 2 there were 645 significant CpG sites between aged and young adult cells (Suppl. File **4a**), 74% of which were hypermethylated in aged vs. young cells. Gene ontology analysis (Suppl. File **4b**) revealed that the DMPs were in genes enriched for ‘cytoskeletal protein binding’ (Suppl. Figure **6c**; Suppl. File **4c**), ‘actin binding’ (Suppl. File **4d**), ‘cytoskeleton’ and ‘development process’ as most significantly enriched, followed by ‘regulation of signaling’ (Suppl. Files **4e, f, g** respectively). KEGG pathway analysis (Suppl. File **4h**) showed enrichment for ‘Insulin secretion’ (Suppl. File **4i**), ‘aldosterone synthesis and secretion’ (Suppl. File **4j**) and ‘cAMP signaling pathway’ (Suppl. Figure **6d**, Suppl. Files **4k**).

It was at 7 d of differentiation that we identified the largest number of DMPs between young and old cells (5,524 DMPs at FDR ≤ 0.05, with a difference/change greater than 2, Suppl. File **5a**). 74% of DMPs were hypermethylated in aged vs. young cells. Gene ontology analysis (Suppl. File **5b**) revealed that DMPs were in genes enriched for ‘developmental process’ (Suppl. Figure **7a**, Suppl. File. **5c**) ‘anatomical structure morphogenesis’ (Suppl. File **5d**), ‘neuron part’, ‘cytoskeletal protein binding’ and ‘cell junction’ (Suppl. File **5 e, f, g**). KEGG analysis (Suppl. File **5h**) identified: ‘Focal adhesion’ (Figure **2d**; Suppl. file **5i**), ‘adherens junction’ (Suppl. File **5j**), ‘regulation of actin cytoskeleton’ (Suppl. File **5k**), ‘cGMP-PKG signaling pathway’ Suppl. File **5l**), ‘rap1 signaling pathway’ (Suppl. File **5m**), as well as the muscle differentiation pathway of ‘PI3K-Akt signaling pathway’ (Suppl. Figure **7b**, Suppl. File **5n**). Among the DMRs at 7 d (Suppl. File **5o**) was a region within *HOXB1*, containing 8 hypermethylated CpGs in aged vs. young cells. We also identified to be hypermethylated in differentiation in aging cells only at 7 days, the gene LY6G5C, this time spanning a slightly larger region Chr6:31650736-31651362, of 627 bp (vs. Chr6:31650736-31651158, 423 bp in the above analysis), confirming that 6 of its CpG’s were hypermethylated at 7 d compared with young adult cells at the same time point. At 10 d, we identified 288 DMPs (FDR ≤ 0.05 with a change greater than 2, Suppl. File **6a**), 89% of which were hypermethylated in aged vs. young cells. Gene ontology analysis (Suppl. File **6b**) revealed that the DMPs were in genes enriched for ‘GDP binding’ (Suppl. Figure **7c**, Suppl. File **6c**), ‘phosphotransferase activity’, ‘positive regulation of antigen receptor’, ‘mesoderm morphogenesis’ and ‘actin binding’ (Suppl. File **6d, e, f, g** respectively). KEGG analysis (Suppl. File **6h**) identified ‘Regulation of actin cytoskeleton’ (Suppl. Figure **7d**; Suppl. File **6i**), ‘ErbB signaling pathway’ (Suppl. File **6j**) and ‘Selenocompound metabolism’ (Suppl. File **6k**).

### Self-organising map (SOM) profiling of aged muscle stem cells confirm a varied methylation profile in aged cells compared with young cells

In order to further analyse temporal dynamics in methylation over the time course of differentiation in aged versus young cells, we conducted SOM profiling analysis of the 2,719 DMPs generated above (Suppl. File **3b** above, those highly significant for ‘age’). SOM analysis averages the methylation values for the group of samples within each condition to identify temporal profile changes in DMPs between aged and young cells over the time-course of differentiation. Aged cells demonstrated more varied DNA methylation changes earlier in the time-course of differentiation (between 72 h and 7 d) compared with young adult cells (Figure **3a**). When looking at aged cells temporal dynamics compared with young cells, out of the 2,719 DMPs in aged cells 1,504 were hypomethylated and 956 hypermethylated at 7 days (confirming the aged cells ‘time’ main effect analysis above). With only a small number of genes demonstrating this altered profile at 7 days in young cells (284 hypomethylated and 110 hypermethylated out of the 2,719 CpG list).

**Figure 3.**
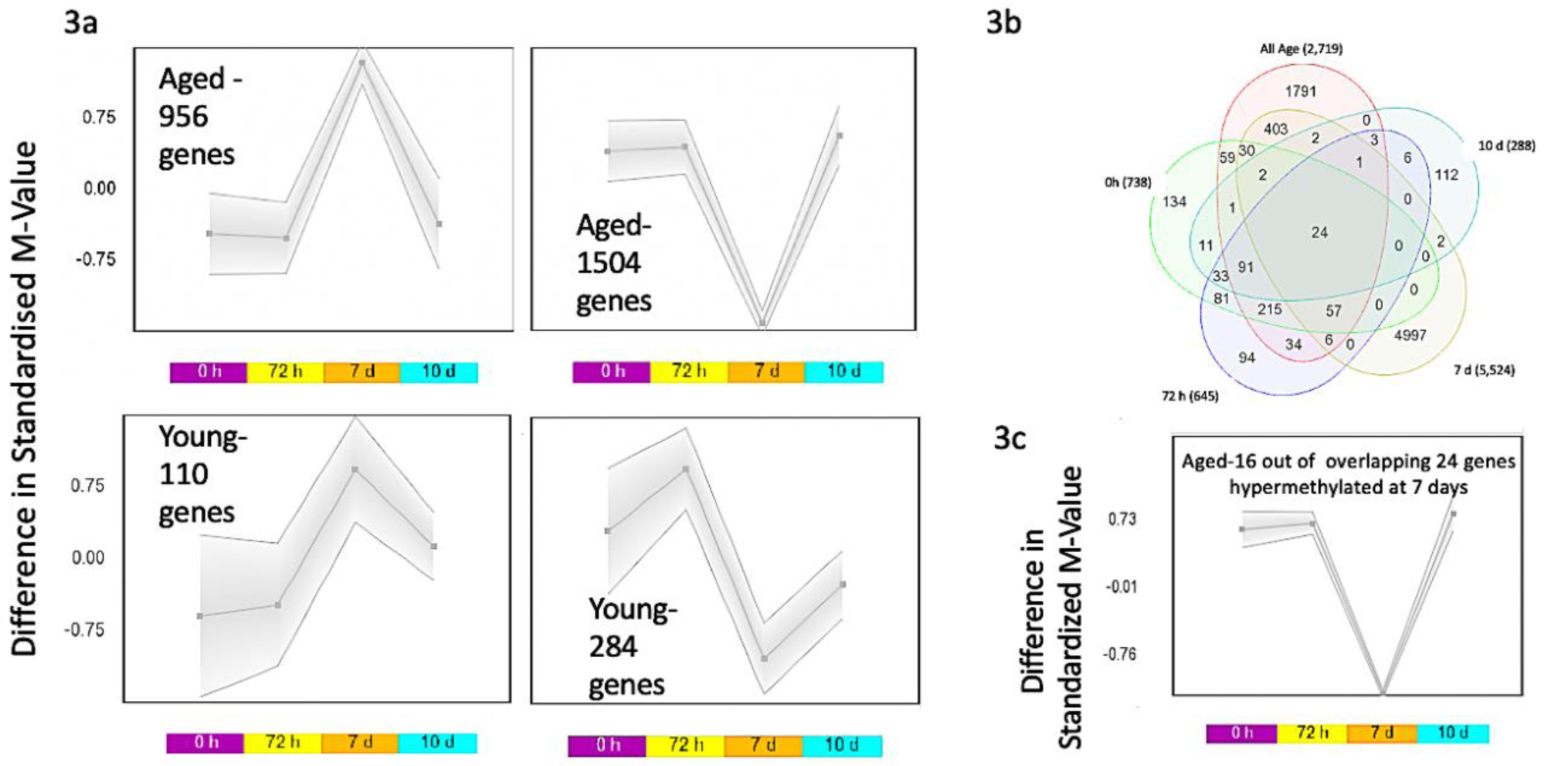
SOM profiling of DNA methylation over the time-course of differentiation in aged versus young adult muscle stem cells. **a**. Demonstrates a larger number of hypomethylated and hypermethylated CpG sites in aged muscle stem cells, particularly at 7 days of differentiation, compared with young adult muscle stem cells. **b**. Venn Diagram analysis depicting the 24 common CpG sites that were altered at every time point of differentiation between aged and young adult muscle stem cells (0, 72 h, 7d and 10 d). **c**. As with the above analysis in 3a, SOM profiling identified that 16 out of these 24 CpG’s also demonstrated the most varied methylation dynamics at 7 days of differentiation in aged cells.

Finally, in order to identify common CpG changes between aged and young cells at each time point, the significantly differentially methylated DMPs from young vs. aged cells across all time were overlapped (2,719 CpG list), with the 0 h aged vs. 0 h young (738 CpG list), 72 h aged vs. young 72 h (645 CpG list), 7 d aged vs. 7 d young (5,524 CpG list), and the 10 d aged vs 10d young (288 CpG list) (Venn diagram, Figure **3b**). There were 24 genes that were identified across tissue and stem cell analysis that were significantly differentially methylated (Suppl. File **7a**). They included 24 CpG’s on 16 genes: TSPAN9, RBM22, UBAP1, CAPZB, ZNF549, MBNL2, RMI1, CHRM5, RAB4A, C19orf21, MOBKL1A, ANAPC11, GAS7, PBX1, ELOVL2, FGGY. Furthermore, generating a SOM profile of temporal change in methylation over the time-course of differentiation in these 24 CpG’s, (Figure **3c**), also demonstrated that the majority of these DMPs in aged cells (16 out of 24 CpG’s) demonstrated varied methylation at 7 days (11 out of 16 CpGs on annotated genes: TSPAN9, RBM22, UBAP1, CHRM5, C19orf21, ELOVL2, MOBKL1A, ANAPC11, PBX1, CAPZB, FGGY; fully detailed in Suppl. File **7a)**.

### Distinguishing differentiation-specific CpG sites in aged cells versus those altered as a consequence of age alone

The data above suggest that aged cells demonstrate hypermethylation versus young adult cells across all stages of differentiation, and that aged cells significantly altered their methylation profiles at 7 days of differentiation. Therefore, we conducted further analysis on the overlap of DMPs within aged cells at 7 d of differentiation (from the ‘time’ analysis above) and those that were changed as a consequence of age at 7 days (aged cells at 7 days versus young cells at 7 d). This enabled the identification of which methylation sites were altered, but also shared in both aged cell differentiation alone and as a consequence of age. Or, alternatively the sites that were simply changed with age and not differentiation process and vice versa. Indeed, overlapping the aged cells 0 h vs. 7 d (1,229 DMP list) with the 7 d young cells significant 5,524 DMP list, there were only 334 (206 hypermethylated, 128 hypomethylated) DMPs that were shared (Suppl. File. **8a**). This suggested that differentiation itself modified only 334 DMPs (out of 1,229) in aged cells that were also changed as a consequence of age (i.e. in aged vs. young cells at 7 days). The remaining 895 DMPs (1,229 - 334 DMPs; Suppl. File **8b**) were differentiation specific to aged cells. Out of this 895 DMP list, an equal number were hypo and hypermethylated (458 hypo and 437 hypermethylated). Therefore, this overlap analysis also confirmed that data above, where over the time-course of aged cell differentiation itself the methylome is both hypo and hyper methylated on a similar number of DMPs, whereas aging alone predominantly hypermethylates (where out of 5,524 changed at 7 d young vs. aged, 4061 were hypermethylated and only 1,463 hypomethylated). This also suggested that the remaining 5,190 (5,524 minus 334 DMP list, equalling 5,190 DMPs including 3,910 hypermethylated vs. 1,280 hypomethylated) were as a consequence of aging alone (Suppl. File **8c**).

### Hypermethylation for a small number of CpG sites is similarly altered in aged muscle cells in-vitro from the in-vivo tissue niche

Tissue and cell PCA plots demonstrated that methylation of even late differentiated cells was vastly different to tissue, suggesting that cell versus tissue samples were comprised of vastly different methylation profiles (Suppl. Figure **8**). Therefore, in order to compare if there were any sites similarly altered in the cells that were also altered in the tissue with age, we overlapped the DMP lists from the tissue and cell analysis described above. Six CpG’s that were identified in the 6,828 significantly differentially methylated CpG list from the aged versus young tissue analysis, as well as highlighted in the 2,719 list ‘age’ cells analysis (Figure **4**). These included: KIF15 (2 CpG’s Cg00702638 & cg24888989), DYRK2 (cg09516963), FHL2 (cg22454769), MRPS33 (cg26792755), ABCA17P (cg02331561). All of these genes were hypermethylated in the tissue analysis as well as the cells. Given that aged cells demonstrated the most varied methylation at 7 days versus young cells. When comparing 7 d aged vs. 7 d young cell list (5,524 list), 4 CpG’s (out of the 6 CpG sites identified above) were also identified in the 6,828 tissue CpG list, including: MGC87042 (cg00394316), C2orf70 (cg23482427), ABCA17P (cg02331561) and cg27209395 (not on an annotated gene). Once more, all of these CpG’s (with the exception of C2orf70, cg23482427) were hypermethylated in the tissue analysis as well as the cells. With ABCA17P (cg02331561) highlighted across all gene lists (**Figure 4**).

**Figure 4.**
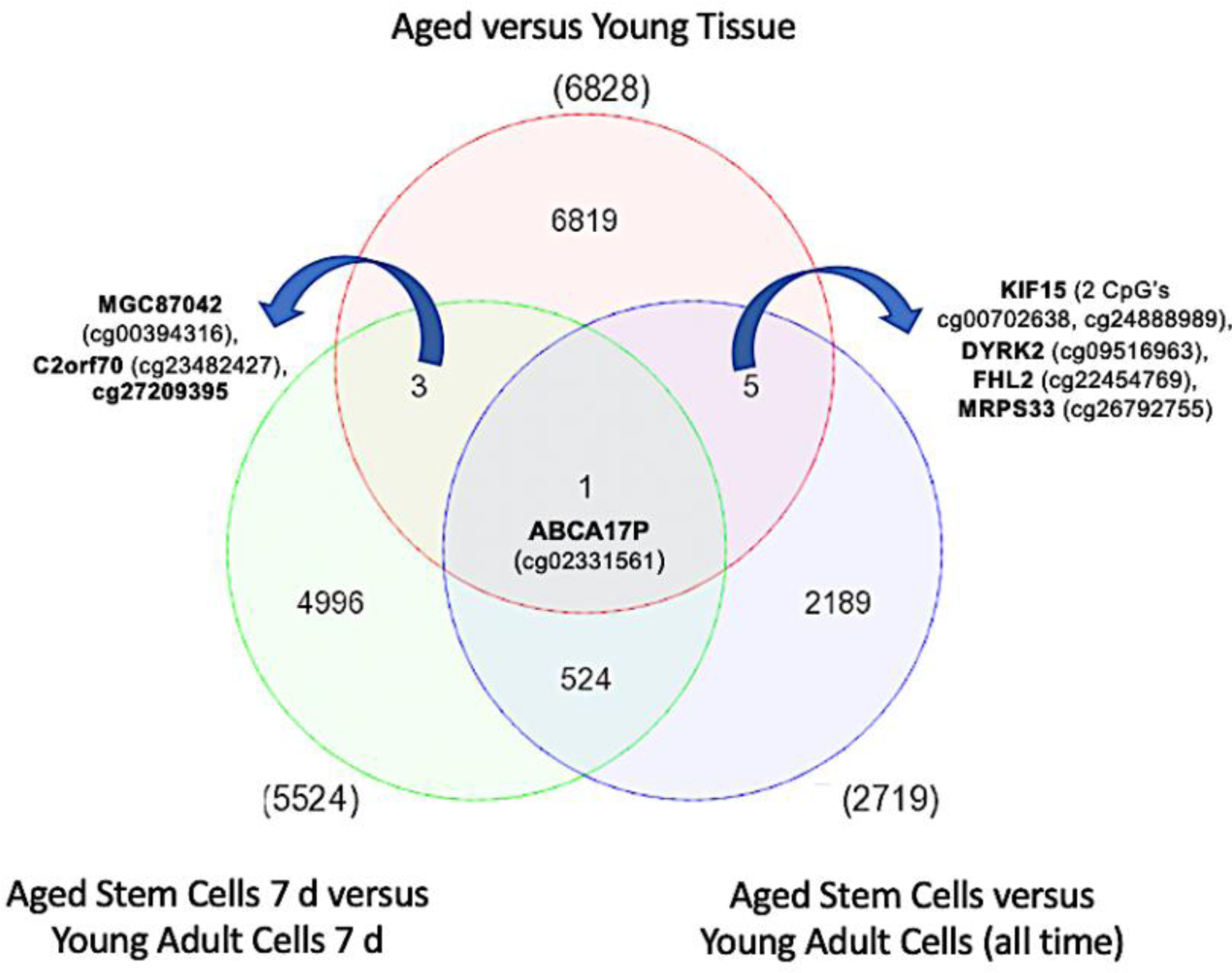
Venn diagram of common overlapping CpG sites significantly differentially methylated between aged skeletal muscle tissue and muscle stem cells. Overlap of the tissue and aged cells identified 6 common CpG sites on genes KIF15 (2 CpG’s Cg00702638 & cg24888989), DYRK2 (cg09516963), FHL2 (cg22454769), MRPS33 (cg26792755) and ABCA17P (cg02331561). Given that aged cells demonstrated the most varied methylation at 7 days versus young cells. When overlapping the 7 d most significantly differentially methylated CpG lists, 4 CpG’s (out of the 6 CpG sites identified above) were also identified, including: MGC87042 (cg00394316), C2orf70 (cg23482427), ABCA17P (cg02331561) and cg27209395 (not on an annotated gene). Once more, all of these CpG’s (with the exception of C2orf70, cg23482427) were hypermethylated in the tissue analysis as well as the stem cells. With ABCA17P (cg02331561) highlighted across all CpG lists.

### Varied methylation in the HOX family of genes in aging skeletal muscle tissue and stem cells

In the above analyses in aged versus young cells (all time points) there were 8 CpG’s with altered methylation within the region of the HOXC10 gene just upstream of HOXC6 and MIR196 (Chr12:54383692-54385621, 1933 bp). Also, on the same chromosome just upstream of the HOXC10 gene (Chr12:54376020-54377868, 1849 bp) within lnRNA HOXC-AS3 there were another 6 CpG’s that were altered in aged cells versus young cells. Similarly, within the 5,524 DMP list at 7 d in aged cells versus 7 d young adult cells (Suppl. File **5o**), the time point most affected by age in the cell analysis, we also identified that a region of the HOXB1, located in Chr17:46607104-46608568 that contained 8 CpG’s that were hypermethylated within the 1465 bp region. Therefore, we further analysed all the HOX gene changes in the significantly differentially methylated aged vs. young tissue (6,828 DMP list) and identified that CpG’s within HOXD10, HOXD9, HOXD8, HOXA3, HOXC9, HOXB1, HOXB3, HOXC-AS2 and HOXC10 were significantly differentially methylated across all analyses, including the aged vs. young tissue (6,828 DMP list; Suppl. File **9a**), aged versus young adult cells (all time) 2,719 DMP list (Suppl. File **9b**) as well as the 7 d aged versus 7 d young cell analysis 5,524 DMP list (Suppl. File **9c**). A Venn diagram analysis depicted the overlap in common gene symbols for these three gene lists (Figure **5a**, Suppl. File **9d**). It was also demonstrated that the majority of these HOX genes were hypermethylated in aged tissue (Figure **5b**; Suppl. File. **9a**). In the cell analysis across all time, these HOX genes also displayed the most varied methylation at 7 days of differentiation in aged cells versus young cells, therefore confirming the varied temporal profile in methylation described above at 7 d was also the case for these HOX genes. Finally, when SOM-profiling these 9 HOX genes by symbol (17 CpGs as some HOX genes contained more than one CpG site), over the time-course of differentiation based on the main effect for ‘age’ 2,719 significantly differentially methylated CpG list. Eight CpG sites across HOX family genes: HOXD8 (cg18448949, cg00035316), HOXB1 (cg04904318, cg02497558, cg22660933, cg10558129), HOXC9 (cg02227188) and HOXB3 (cg09952002) were hypermethylated at 7 days, whereas and 7 CpGs across HOXC10 (cg20402783, cg20403938, cg16898193, cg16791250), HOXB3 (cg23014425, cg04800503), HOXC-AS2 (cg09605287) were hypomethylated at 7 days (Figure **5c;** Suppl. File **9e**). This meant distinct genes were hypermethylated (HOXD8, HOXC9, HOXB1) and hypomethylated (HOXC10, HOXC-AS2) at 7 days in aged cells versus young adult cells, except for one of these genes, HOXB3, that contained 1 CpG that was hypermethylated versus 2 that were hypomethylated.

**Figure 5.**
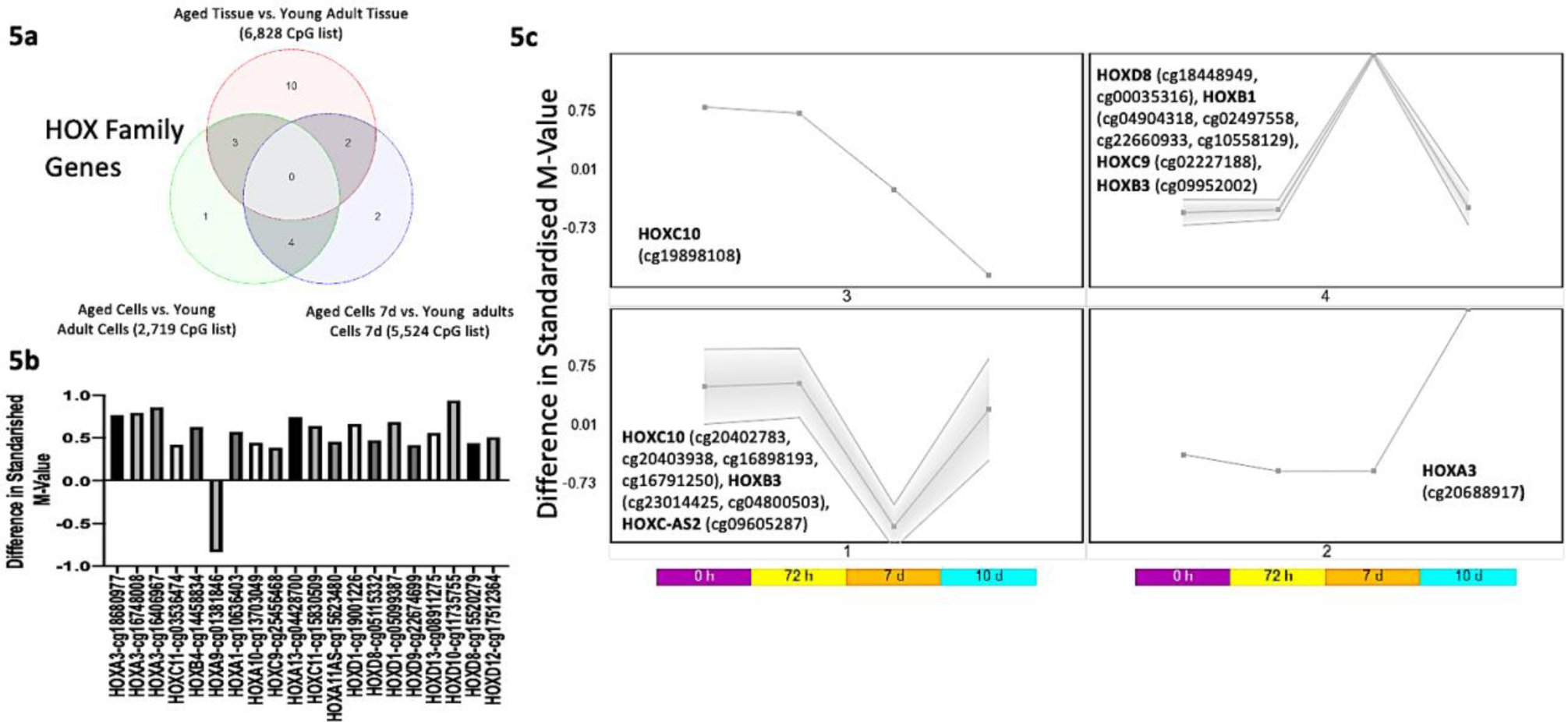
HOX family of genes and their DNA methylation in aged tissue and muscle stem cells. **a**. Venn diagram identifying 9 commonly differentially methylated HOX genes: including HOXD10, HOXD9, HOXD8, HOXA3, HOXC9, HOXB1, HOXB3, HOXC-AS2 and HOXC10 (note this Venn diagram analysis is by ‘gene symbol’ not ‘probe cg’ (CpG site), as some HOX genes also had more than 1 CpG per gene symbol, full CpG lists are located in Suppl. Figure **9 a**,**b**,**c**,**d**). **b**. All HOX family genes by CpG site (cg probe) differentially methylated in aged compared with young skeletal muscle tissue, predominantly all demonstrating hypermethylation. **c**. SOM profiling depicting the temporal regulation of DNA methylation in aged muscle stem cells as they differentiate in CpGs located amongst the HOX family of genes (depicting the 9 HOX genes altered, by gene symbol, in both the tissue and cells). The majority of these HOX CpG sites were differentially methylated at 7 days of differentiation in the aged cells.

We next analysed the gene expression of the HOX genes that changed at the methylation level in both the tissue and cell analysis (by gene symbol-HOXD8, HOXA3, HOXC9, HOXB1, HOXB3, HOXC-AS2 and HOXC10). Interestingly, in aged tissue, despite displaying hypermethylation in all these HOX genes versus young tissue, the genes were not suppressed at the gene expression level, which may have been expected, yet all elevated (Suppl. Figure **9**). However, in the aged cells when analysing these genes at the expression level at 7 days of differentiation (including HOXD8, HOXA3, HOXC9, HOXB1, HOXB3, HOXC-AS2 and HOXC10, as well as HOXC-AS3 identified in the above in the cell analysis only) (Figure **6**), we identified that there was significantly reduced gene expression in gene HOXB1 (Figure **6**), that was inversely related with increased HOXB1 methylation. Where in the 5,524 DMP list at 7 d in aged cells versus 7 d young adult cells (Suppl. File **5o**), we previously identified that a region of the HOXB1 located in Chr17:46607104-46608568, contained 8 CpG’s that were hypermethylated, as well as HOXB1 Cg’s: cg04904318, cg02497558, cg22660933, cg10558129 being identified as hypermethylated in the 2,719 significant main effect for ‘age’ cell CpG list above (Suppl. File **3b**). There was also significantly increased gene expression for HOXC-AS3, with this gene identified earlier in the analysis as being having reduced (hypo)methylation. Indeed, hypomethylation occurred in 6 CpG’s in a region upstream of the HOXC10 gene (Chr12:54376020-54377868, 1849 bp) within the HOXC-AS3 gene. Interestingly, HOXC10 also demonstrated an average increase in gene expression, however, it was not statistically significant (Figure **6**). Finally, HOXA3 also demonstrated significantly reduced expression (Figure **6**) with corresponding hypermethylation (at 10 not 7 days of differentiation) in aged cells (see Figure **5c**). Overall, HOXB1, HOXC-AS3 and HOXA3 demonstrated an inverse relationship with CpG methylation and gene expression in aged versus young cells.

**Figure 6.**
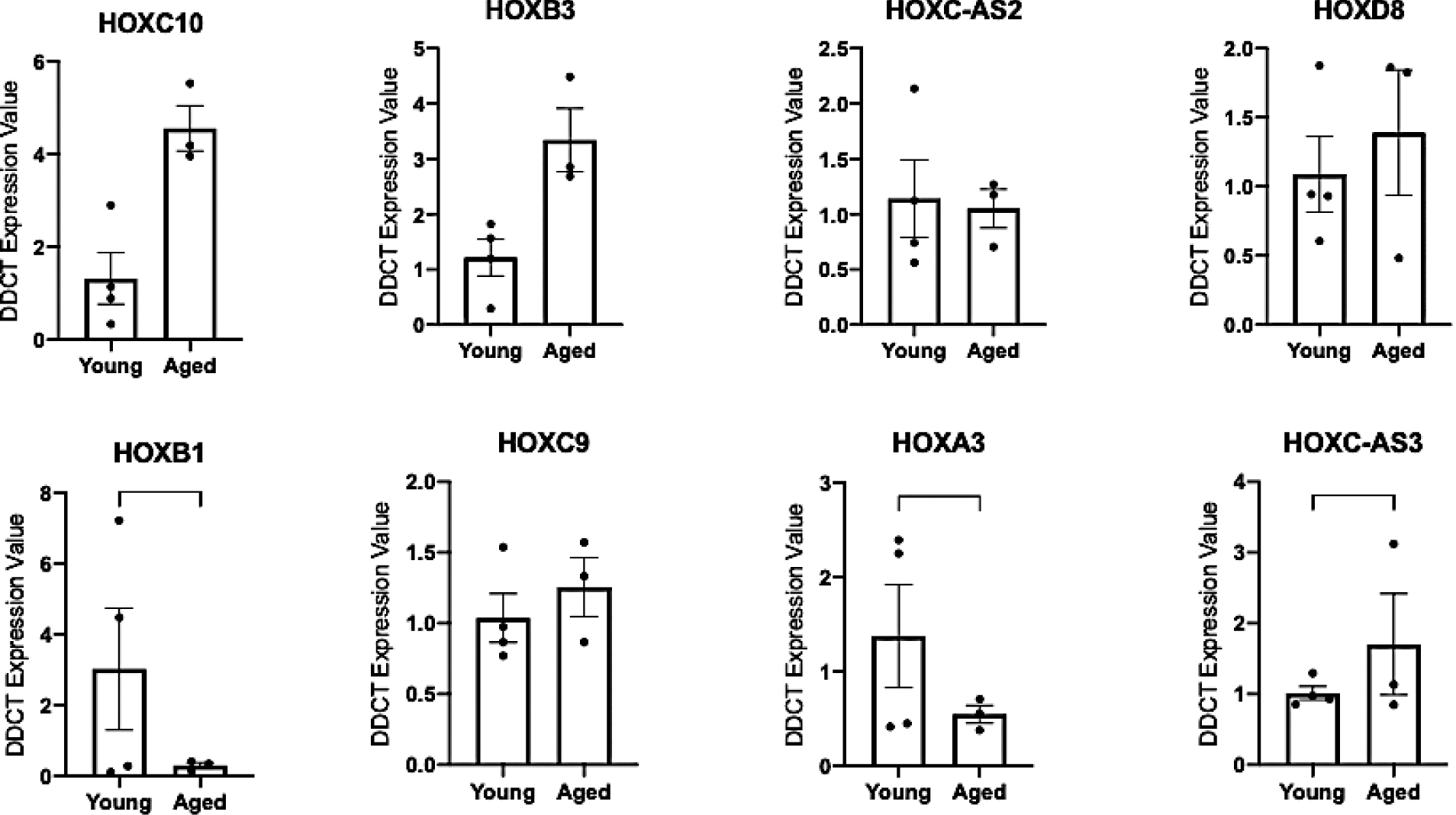
Gene expression of the HOX family of genes in aged compared to young muscle stem cells at 7 days of differentiation.

### Effect of physical activity on methylation status of HOX family genes

Next, given that aging generally hypermethylated the genome, we tested the hypothesis that increasing physical activity may oppositely regulate DNA methylation and be associated with increasing hypomethylation in these HOX genes. We thus performed a multiple regression analysis using methylation data and the level of physical activity of 30 endurance-trained men. As in the above analysis, we also found that CpG-sites associated with physical activity (P<0.05) were significantly enriched with HOX genes (137 of 1219 CpG-sites, Fisher’s exact test OR=1.7, P=2.3*10-8, Suppl. File **10a)**. Where we determined that highly active men had hypomethylated HOXB1 (cg10558129, P=5.2*10^−4^), HOXA3 (cg16406967, P=0.03), HOXD12 (cg17512364, P=0.008) and HOXC4 gene (cg13826247, P=0.014) compared to less active men (adjusted for age and muscle fiber composition) on the same sites identified above in the aging data. Furthermore, we identified hypomethylation of HOXA3 on several additional sites (cg21134232, P=0.0014, cg25768734, P=0.0002; cg27539480, P=0.039; cg00431187, P=0.026; cg03483713, P=0.045; cg15982700, P=0.026; cg23806243, P=0.013). Full methylome analysis of the physical activity dataset can be found in Suppl. File **10b**, with details of all HOX sites in Suppl. File **10a**). Given that we identified the opposite trend in aged muscle and cells, particularly *HOXB1* and *HOXA3* that were hypermethylated in aged tissue and stem cells, and with reduced gene expression in aged cells. These findings suggest that increasing levels of physical activity are associated with increasing reductions in methylation (hypomethylation) in these HOX genes compared to age-related changes that are associated with increasing methylation (hypermethylation).

## Discussion

In the present study we first aimed to investigate the methylome in aged skeletal muscle tissue and differentiating primary muscle stem cells compared with young adults, in order to identify important epigenetically regulated genes in both aging skeletal muscle tissue and muscle derived stem cells. As with previous studies ^18, 27^, and by using more recent, higher coverage array technology, we identified that aged skeletal muscle tissue demonstrated a considerably hypermethylated profile compared with young adult tissue. We also demonstrated that these hypermethylated profiles in aged tissue were enriched in gene ontology pathways including, ‘regulation of muscle system process’ and KEGG pathways ‘pathways in cancer’, a pathway that incorporates previously well described molecular pathways in the regulation of skeletal muscle such as; focal adhesion, MAPK signaling, PI3K-Akt-mTOR signaling, p53 signaling, Jak-STAT signaling, TGF-beta and Notch signaling, as well as the other significantly enriched pathways of ‘rap1 signaling’, ‘axon guidance’, and ‘hippo signaling’. This was also the first study to profile DNA methylation over the entire time-course of skeletal muscle differentiation (0, 72 h, 7 and 10 d) using the highest coverage 850K methylation assays. In primary cell cultures, isolated from aged and young adults matched for the proportion of myogenic cells, we identified that aged muscle stem cells also demonstrated hypermethylated profiles versus young adult cells. This hypermethylation was enriched in: ‘axon guidance’, ‘adherens junction’ and ‘calcium signaling’ pathways. Furthermore, we identified that the process of cellular differentiation itself did not significantly affect DNA methylation in young cells, however aged cells demonstrated varied methylation profiles particularly at 7 d of differentiation in GO terms: ‘regulation of localisation’, ‘regulation of cell communication’, and ‘regulation of signaling’. Also, in the majority of different CpG sites that were altered during the process of differentiation, aged cells demonstrated significantly hypermethylated profiles during differentiation when compared with young adult cells. Again, specifically at 7 d of differentiation including CpG’s located within: ‘focal adhesion’, ‘adherens junction’ and ‘regulation of actin cytoskeleton’ pathways, as well as the well-known muscle differentiation pathway, ‘PI3K-AKT signalling’. This corresponded with reductions in myoD and myogenin, and delayed increases in myogenin gene expression in aged compared with young adult cells.

We were also able to identify that a small number of CpG sites hypermethylated in aged tissue were also hypermethylated in aged cells, with CpG’s located on genes: KIF15, DYRK2, FHL2, MRPS33, ABCA17P. This suggested, that perhaps these CpG’s retained their methylation status *in-vitro* after being isolated from the *in-vivo* niche. However, it is worth noting that this was a very small number of CpG’s compared with the thousands of CpG sites significantly differentially methylated in the aged tissue versus young adult tissue, and those also observed to be significantly different in the aged versus young muscle cells. This was suggestive that the majority of the hypermethylated CpG’s observed at the aged tissue level were generally not retained on the same CpG sites in the isolated aged muscle stem cells *in-vitro*. Also, PCA plots of muscle tissue versus isolated muscle cells demonstrated vastly different profiles, suggesting that isolated cells, even late differentiated muscle cells, are perhaps really quite different epigenetic representations of contrasting cellular entities versus muscle tissue. This also perhaps indicates that hypermethylation of DNA within myonuclei (the predominant source of DNA in the tissue samples) maybe therefore unique to that observed in the isolated aged muscle progenitor cells in combination with other muscle derived cell populations. Indeed, retention of methylation during aging has been previously observed in artificially aged muscle cells ^19^. In skeletal muscle tissue, retention of DNA methylation has been observed after skeletal muscle growth, even after a subsequent period of muscle loss, suggesting an epigenetic memory at the DNA methylation level ^21, 22, 49^. However, the relative contribution of methylation from myonuclei or satellite cells (or other resident cell types in muscle tissue) to this epigenetic memory effect, and how long these retained profiles can last (e.g. past 22 weeks in the Seaborne *et al*., 2018 study) has not been determined. However, interestingly based on the present studies data, we could hypothesise that myonuclear hypermethylation in the tissue with age is quite different to the hypermethylation observed in muscle progenitor cells as they differentiate. Perhaps, suggesting that environmental stimuli and aging could affect methylation profiles in the myonuclei differently than those in satellite cells. A hypothesis that requires further investigation, perhaps using single-cell DNA/RNA analysis of the different cell populations resident in cells derived from skeletal muscle tissue biopsies compared with cultured cells. Finally, it is important to mention that the human derived muscle cells were comprised of both muscle and non-muscle cells in an attempt to capture the changes across the skeletal muscle ‘milieu’ of cell types. Therefore, methylation profiles reflect both the differentiation of muscle cells but also expansion predominantly fibrogenic and a minority of other adherent muscle derived cells. As suggested above, with respect to specific methylation changes in myonuclei versus satellite cells, it will be important in future studies to undertake single-cell DNA/RNA analysis to identify the specific contribution of different cell types to the methylation profiles observed in aged cells. Indeed, exciting recent work suggests different methylation profiles from myonuclear versus interstitial cells in response to a hypertrophic stimulus in skeletal muscle ^62^.

Importantly, in both tissue and stem cell analysis, we also identified that the homeobox (HOX) family of genes were significantly enriched in differentially methylated region analysis, showing several (e.g. 6-8) CpGs to be methylated within chromosomal regions on these genes in aged compared with young adults. In particular, we identified: HOXC10 (just upstream of HOXC6) and HOXB1 as having several CpGs differentially methylated. Therefore, closer analysis of all HOX gene associated CpG changes across both tissue and cell differentiation data identified that CpG’s located within: HOXD10, HOXD9, HOXD8, HOXA3, HOXC9, HOXB1, HOXB3, HOXC-AS2 and HOXC10 were all significantly differentially methylated across these analyses. In aged tissue the majority of these HOX genes were hypermethylated. In the cell analysis, these HOX genes displayed the most varied methylation at 7 days of differentiation in aged versus young cells. Furthermore, distinct HOX genes were hypermethylated (HOXD8, HOXC9, HOXB1) and hypomethylated (HOXC10, HOXC-AS2) at 7 days in aged cells versus young adult cells. Gene expression analysis also demonstrated an inverse relationship with DNA methylation. Where hypermethylation of HOXB1 and HOXA3 was associated with reduced gene expression, and hypomethylation of HOXC-AS3 associated with increased gene expression in aged versus young cells.

HOX genes are evolutionary conserved members of the homeobox superfamily, with 39 HOX genes found in the mammalian genome. They are ‘homeotic’ genes coding for transcription factors, with a fundamental role in the determination of cellular identity. They were first shown to be important in embryogenesis in drosophila melanogaster (fruit fly) ^63^. HOX genes have also been reported to alter with age in non-muscle tissues ^64, 65^. In muscle they have been described to morphologically identify the hindlimb tissues during development ^66, 67, 68, 69^, but have also been demonstrated to be activated in satellite cells ^70, 71, 72^, and as markers of hindlimb derived myoblasts ^70^. In particular HOXC10, demonstrated 8 CpG’s, just upstream of HOXC6 and miR-196; Chr12:54383692-54385621, 1933 bp that all demonstrated a hypomethylated signature in aged versus young muscle stem cells. Where miR-196 has previously been demonstrated to regulate HOX gene expression in adipose tissue ^73^. There were also 4 CpG sites hypomethylated, particularly at 7 days of differentiation in aged versus young cells. Indeed, HOXC10 has been identified to determine hindlimb identity ^66, 68^. Together with HOXC10 hypomethylation, we also demonstrated average (yet not significant) increases in HOXC10 gene expression at 7 days in aged cells versus young cells. Counterintuitively to our data, previously HOXC10 upregulation has been associated with satellite cell activation in skeletal muscle in response to Roux-en-Y gastric bypass (RYGB) surgery ^74^, as well as being a marker for hindlimb specific satellite cells. Interestingly, there was lower expression of HOXC10 observed in exercised rats ^75^, which is perhaps more intuitive with the data provided here, where aged cells demonstrated an increase. However, HOXC10 requires more experimentation to define its mechanistic role in aging skeletal muscle, with HOXC10 and physical exercise being discussed below. Interestingly, the hypomethylation of the HOXC10 (Chr12:54383692-54385621, 1933 bp) occurred in 8 CpG’s on the same chromosome just upstream of the HOXC10 gene (Chr12:54376020-54377868, 1849 bp), within the lncRNA HOXC-AS3, where there were another 6 CpG’s that were hypomethylated in aged cells versus young cells (See Suppl. Figure **10** for visualisation of HOXC10 and HOXC-AS3 and their methylation and genomic location). HOXC-AS3 is a long coding RNA (lcRNA) with currently no known data in skeletal muscle, with some literature in cancer and mesenchymal stem cell (MSC) fields ^76^. Indeed, MSC’s administered with silencing HOXC-AS3 prevented bone loss by enhancing HOXC10 expression ^77^, and HOXC-AS3 upregulation has also been linked with aggressive cancer ^78^. Interestingly, we were able to identify that together with associated hypomethylation, in a region close to the lcRNA HOXC-AS3, there was significantly increased gene expression of HOXC-AS3 (and an average, yet not significant increase in HOXC10 gene expression) in aged muscle cells at 7 days of differentiation. Given the data above in bone and cancer, HOXC-AS3 upregulation appears to be pro-growth and linked with expression of HOXC10, therefore their increase in the current study maybe hypothesised to be a co-operative and compensatory drive to maintain aged muscle. However, this hypothesis is speculative, and more work needs to be conducted as to the role of HOXC10 and HOXC-AS3 and their potential cooperative mechanisms of action in aged skeletal muscle.

We also identified that HOXB1 was hypermethylated with increased gene expression in aged cells at 7 days. HOXB1 has been demonstrated to be hypermethylated in inflamed muscle of children with Juvenile Dermatomyositis (JDM) ^79^. This is interesting given aged skeletal muscle is known to be chronically inflamed ^80, 81^, where we also demonstrate this hypermethylated profile in aged cells. HOXA3 was hypermethylated with reduced gene expression in aged cells. However, there is currently little to no work on this gene in skeletal muscle ^69^ and therefore this requires future investigation. It is however worth noting that gene expression, in the aged tissue in particular, was not expected given the methylation data. Where aged tissue demonstrated an increase in gene expression of HOX genes with the majority demonstrating hypermethylation. This may be due one of the elderly patients’ donors demonstrated much greater expression versus all the other donors for some of the HOX genes. Therefore, in order to confirm this HOX gene expression would require confirmation in a larger population of aged patients. Other limitations to the study include a mixed cohort of male and female aged individuals compared to the young adult group being all male. PCA analysis of male and female aged samples did not show differences between methylation profiles and we also removed sex (X and Y) chromosome probes from the analysis. Furthermore, the muscle tissue and cells were isolated from two sites, the gluteus medius and vastus lateralis in the aged samples compared with only the vastus lateralis in the young adult samples. This is important to note as the gluteus medius and vastus lateralis have different fibre type proportions. Therefore, the data should be viewed with that caveat in mind. Despite this, the changes in methylation of the sites observed were identified across samples from both biopsy sites, suggesting that these changes occurred in aged individuals across muscle types.

Finally, with aging evoking a hypermethylated signature in tissue and aged muscle derived stem cells, it was also interesting to speculate that physical exercise, that has been shown to hypomethylate the genome ^20, 21, 22^, could therefore be ‘anti-ageing’ at the epigenetic level. Indeed, this hypothesis was supported indirectly in the present study and by previous literature, where the aged tissue analysis in the present study identified the top significantly enriched KEGG pathway as, ‘pathways in cancer’. A pathway that incorporates well known pathways important in skeletal muscle, including: Focal adhesion, MAPK, PI3K-Akt, mTOR, p53 signaling, Jak-STAT, TGF-beta and Notch signaling. Where, this pathway was also the top enriched hypomethylated pathway after acute and chronic resistance exercise ^22^. While, perhaps the significance of these larger pathways such as ‘pathways in cancer’ can be inflated in methylation analysis ^82^, and therefore should be viewed with some caution. This data perhaps suggests that exercise (resistance exercise) could perhaps reverse the hypermethylated profiles in these pathways in aged muscle. Therefore, in the present study, given that we identified the HOX family of genes to be extensively differentially methylated in aged tissue and stem cells, we went on to determine that increasing physical activity levels (endurance exercise) in healthy young adults was associated with larger reductions in HOXB1 and HOXA3 methylation (hypomethylation). This was opposite to the changes we observed with age in both muscle tissue and stem cells, that demonstrated increased hypermethylation with age. This also provided evidence to suggest that increased physical activity could perhaps reverse the age-related epigenetic changes in the HOX genes. Importantly, HOXA3 has been shown to be hypomethylated on multiple sites after resistance exercise training, with retention of hypomethylation for site (cg12434681) during detraining into retraining ^21^. Suggesting, that this HOX gene is also hypomethylated with exercise training and possesses an epigenetic memory from earlier exercise training as previously described by our group ^60^. Unpublished work by our group also suggests that acute sprint exercise in human muscle can hypomethylated the same HOXA3 CpG site (cg00431187) that was oppositely hypermethylated in aging but demonstrated hypomethylation in individuals that have higher physical activity level. However, more research into the effect of exercise in an aged population and the changes in HOX methylation status will be required in the future to confirm these findings. Our overarching main results from this study are summarised schematically in Figure 7.

**Figure 7.**
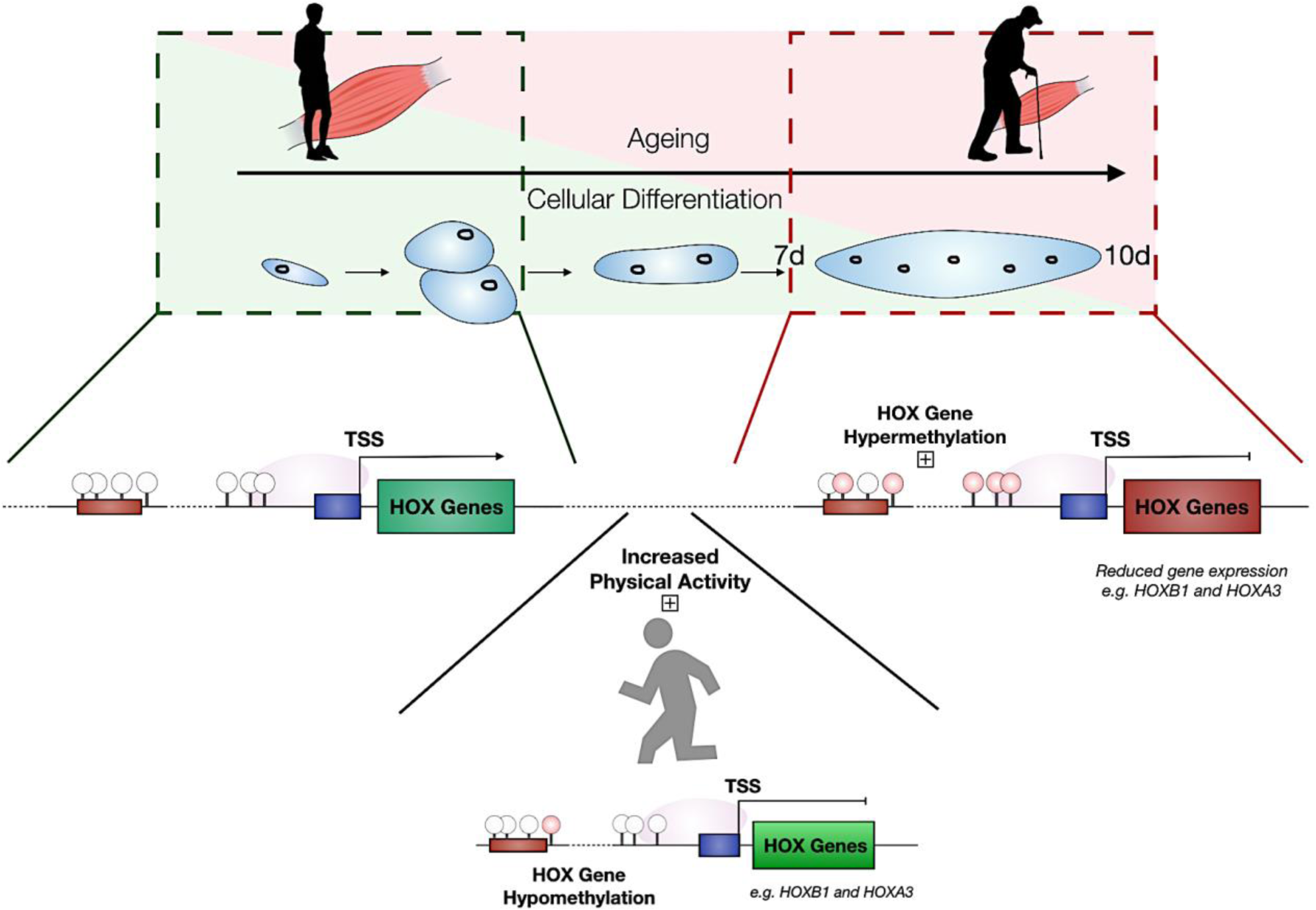
Schematic of the overarching main results. Hypermethylation of the genome and of the HOX genes in aging skeletal muscle tissue (RED box) compared with young adult tissue (GREEN BOX). Dysregulation of HOX genes in aged cells particularly at 7 days of differentiation in muscle derived cells. Inverse methylation (hypermethylation-red circles) with gene expression (reduced-red rectangle) in aging cells particularly for *HOXB1* and *HOXA3*. Increased physical activity levels are associated with the hypomethylation (fewer red circles more white circles) of the same two genes, *HOXB1* and *HOXA3*.

## Conclusion

Overall, for the first time, we demonstrate that altered methylation of a large number of the HOX genes are epigenetically differentially regulated in aged human skeletal muscle tissue and during impaired differentiation in aged muscle stem cells. In particular HOXB1, HOXA3 and HOXC-AS3 (and to a certain extent, HOXC10) also demonstrated significantly inversed changes in gene expression in aged cells. Finally, that increased physical activity may help prevent the age-related epigenetic changes observed in these HOX genes

## Acknowledgments and funding

These data were supported by a North Staffordshire Medical Institute (NMSI) grant awarded to Adam P. Sharples (PI), Daniel Turner, Mark Kitchen and Ian Dos-Remedios (Co-I’s). This work was also supported by the UK’s Engineering and Physical Sciences Research Council (EPSRC) and the UK Medical Research Council’s (MRC) centre for doctoral training, via a studentship awarded to the joint first author Piotr Gorski in the group of Adam P. Sharples (PI). Funds from Keele University and Liverpool John Moores University, UK and

The Norwegian School of Sport Sciences, Oslo, Norway also supported the PhD work by Daniel Turner & Piotr Gorski in the group of Adam P. Sharples. Philipp Baumert received a fully-funded Liverpool John Moores PhD scholarship. Mohd Firdaus Maasar received a PhD studentship via the Malaysian government agency: Majlis Amanah Rakyat (MARA) via Barry Drust, Adam P. Sharples and Dr. Andrew Hulten. The physical activity and DNA methylation study was supported in part by grant from the Russian Science Foundation (Grant No. 17-15-01436: “Comprehensive analysis of the contribution of genetic, epigenetic and environmental factors in the individual variability of the composition of human muscle fibers”; DNA sample collection, genotyping, epigenetic analysis and muscle fibre typing of Russian subjects).

## Declaration

All authors declare no conflicts of interest.

