## Supplementary material for "DNA methylation across the genome in aged human skeletal muscle tissue and stem cells: The role of HOX genes and physical activity": Suppl. Figure 1

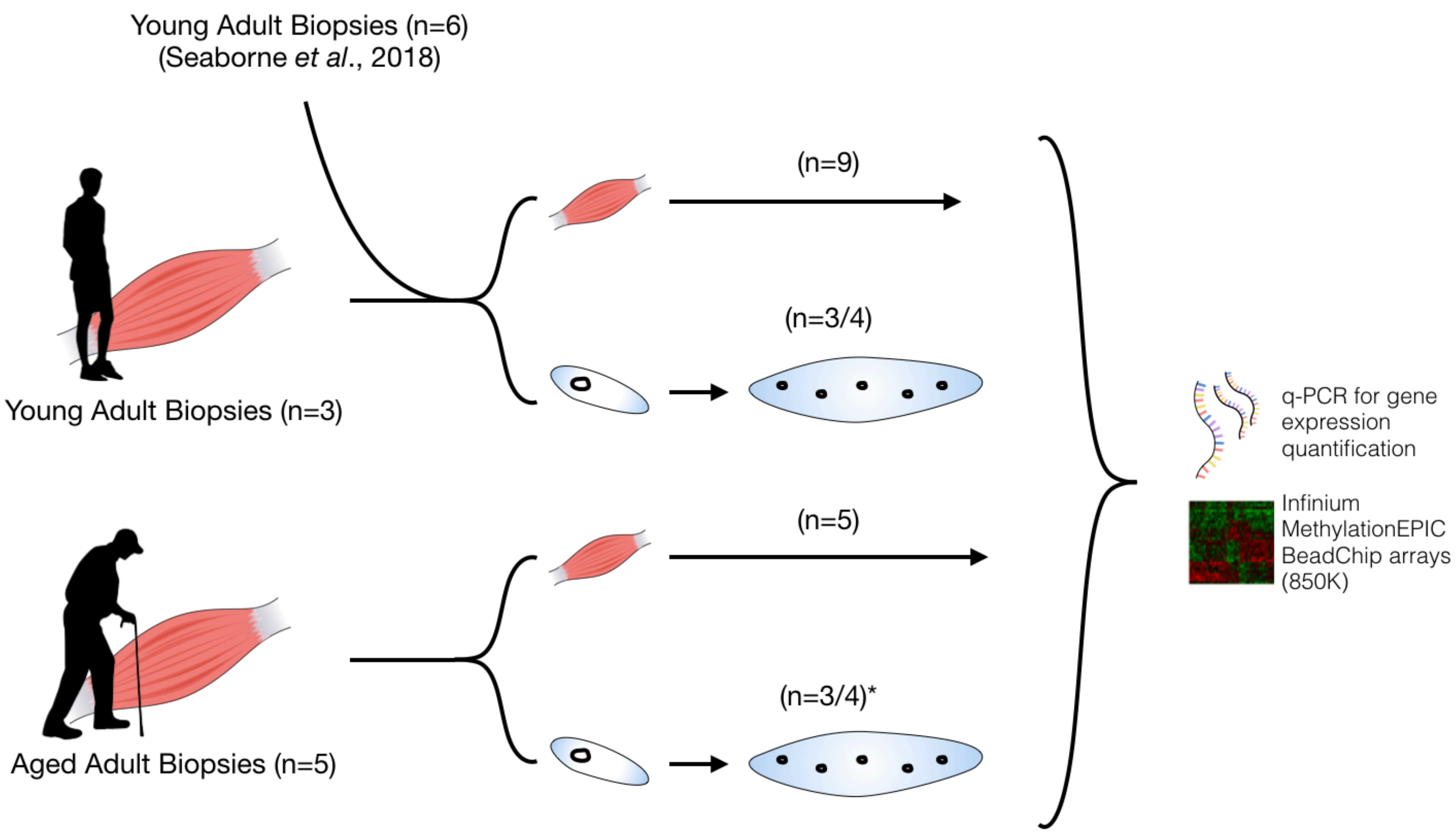


**Suppl. Figure 1.** Schematic of experimental approach (* 10 d Young cells n = 2).

**Title:** DNA methylation across the genome in aged human skeletal muscle tissue and stem cells: The role of HOX genes and physical activity

**Authors:** Turner DC, Gorski PP, Maasar MF, Seaborne RA, Baumert P, Brown AD, Kitchen MO, Erskine RM, Dos-Remedios I, Voisin S, Eynon N, Sultanov RI, Borisov OV, Larin AK, Semenova EA, Popov DV, Generozov EV, Stewart CE, Drust B, Owens DJ, Ahmetov II, Sharples AP.
