## Supplementary material for "DNA methylation across the genome in aged human skeletal muscle tissue and stem cells: The role of HOX genes and physical activity": Suppl. Figure 2

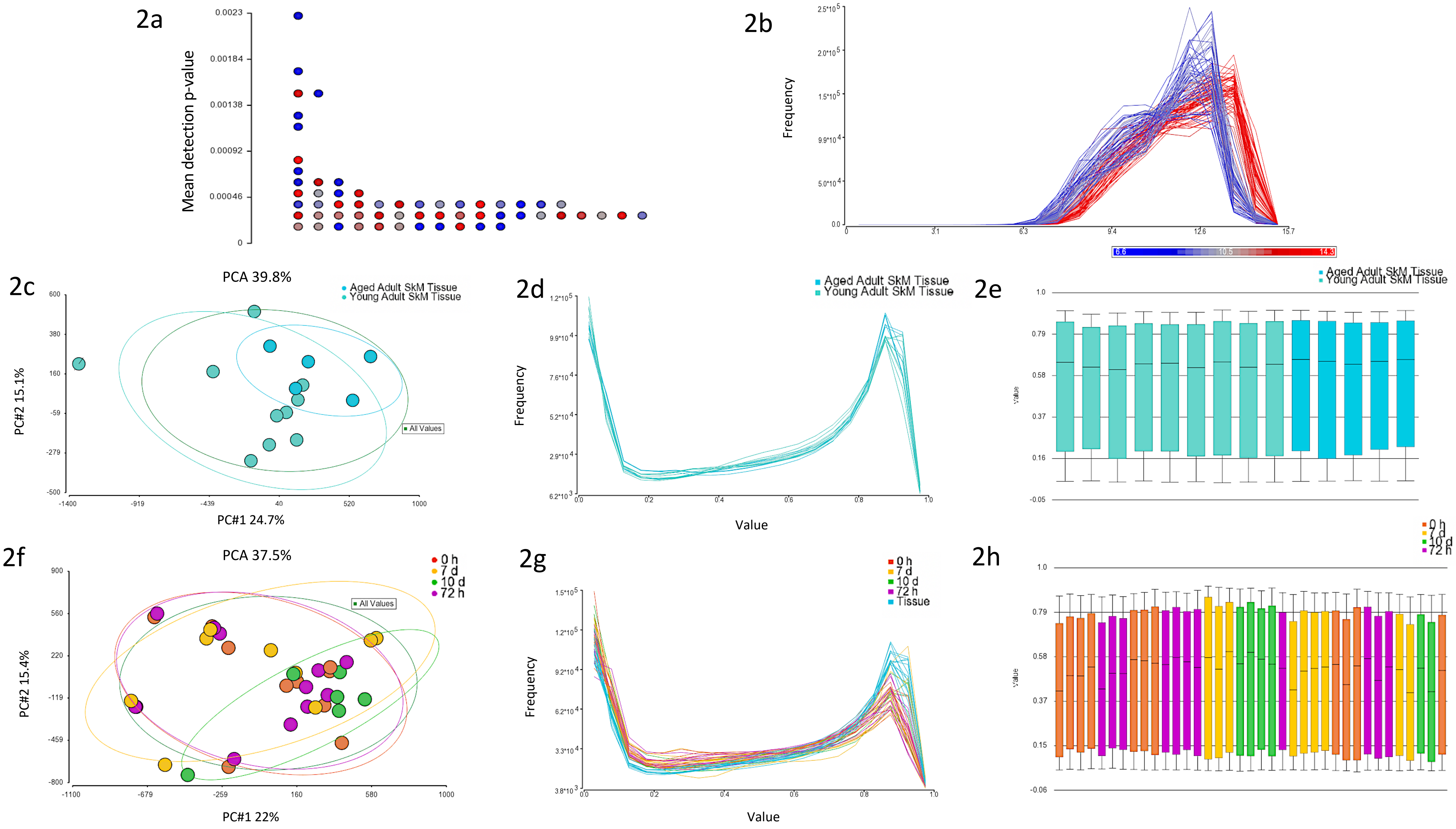
2a
Mean detection p-value
0.0023
0.00184
0.00138
0.00092
0.00046
0
2b
Frequency
2.5*10^5
2.0*10^5
1.5*10^5
9.9*10^4
5.0*10^4
0.0
0
3.1
6.3
9.4
12.6
15.7
6.6
10.5
14.3
2c
PCA 39.8%
Aged Adult SkM Tissue
Young Adult SkM Tissue
All Values
PC#2 15.1%
PC#1 24.7%
600
380
160
-59
-279
-500
-1400
-919
-439
40
520
1000
2d
Aged Adult SkM Tissue
Young Adult SkM Tissue
Frequency
Value
1.2*10^5
9.9*10^4
7.6*10^4
5.2*10^4
2.9*10^4
6.2*10^3
0.0
0.2
0.4
0.6
0.8
1.0
2e
Aged Adult SkM Tissue
Young Adult SkM Tissue
value
1.0
0.79
0.58
0.37
0.16
-0.05
2f
PCA 37.5%
0 h
7 d
10 d
72 h
All Values
PC#2 15.4%
PC#1 22%
900
560
220
-119
-459
-800
-1100
-679
-259
160
580
1000
2g
0 h
7 d
10 d
72 h
Tissue
Frequency
Value
1.5*10^5
1.2*10^5
9.2*10^4
6.2*10^4
3.3*10^4
3.8*10^3
0.0
0.2
0.4
0.6
0.8
1.0
2h
0 h
7 d
10 d
72 h
value
1.0
0.79
0.58
0.37
0.15
-0.06

**Suppl. Figure 2. QA and QC of Illumina EPIC 850K DNA methylation arrays. 2a.** Average detection p-value of probes per sample. The highest average detection p value was 0.0023, well below the 0.01 cut-off. Each dot represents the average detection p-value for each sample. **2b.** Density plots of the raw intensities/signals of the probes in each sample, demonstrated all methylated and unmethylated signals were over 11.5 (mean was 11.52 and median was 11.8), and the the difference between median methylation and median unmethylated signal was 0.56. **2c:** PCA aged vs. young adult tissue, with outlier outside 2SD's of ellipsoids highlighted (line strikethrough- single sample) that was subsequently removed from the analysis. **2d.** Frequency plot by lines of aged vs. young adult tissue with outlier identified in 2c above, removed. **2e.** Box and whisker plots of aged vs. young adult tissue. **2f.** PCA of aged and young adult muscle stem cells over the time-course of differentiation, demonstrating no outliers. **2g.** Frequency plot by lines of aged vs. young adult muscle stem cells demonstrating no outliers in frequency distribution compared with other samples. **2h.** Box and whisker plots of aged vs. young adult muscle stem cells.

**Title:** DNA methylation across the genome in aged human skeletal muscle tissue and stem cells: The role of HOX genes and physical activity

**Authors:** Turner DC, Gorski PP, Maasar MF, Seaborne RA, Baumert P, Brown AD, Kitchen MO, Erskine RM, Dos-Remedios I, Voisin S, Eynon N, Sultanov RI, Borisov OV, Larin AK, Semenova EA, Popov DV, Generozov EV, Stewart CE, Drust B, Owens DJ, Ahmetov II, Sharples AP.
