## Supplementary material for "DNA methylation across the genome in aged human skeletal muscle tissue and stem cells: The role of HOX genes and physical activity": Suppl. Figure 4

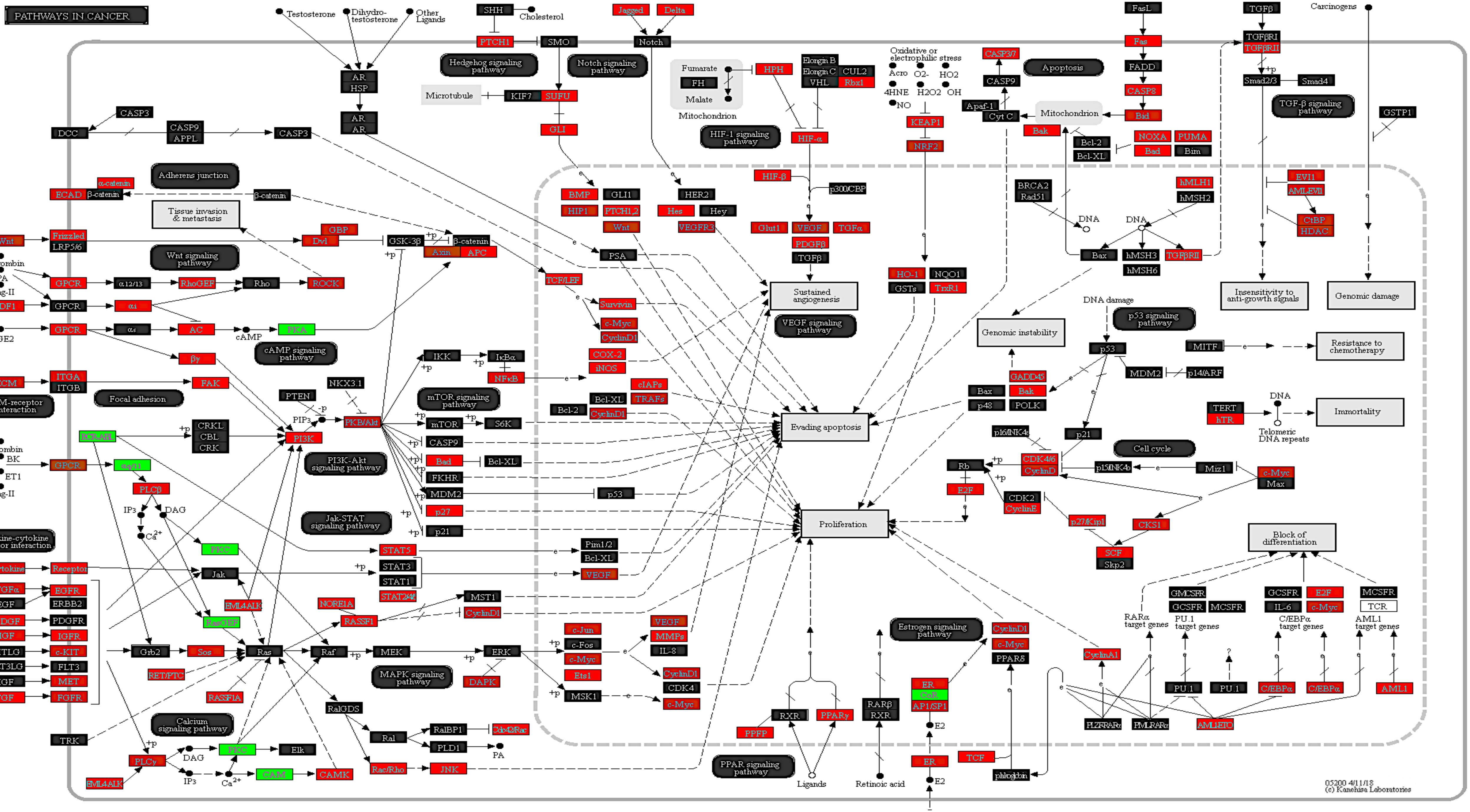

PATHWAYS IN CANCER
Testosterone
Dihydro-testosterone
Other Ligands
AR
HSP
SHH
Cholesterol
PTCH1
SMO
Hedgehog signaling pathway
Microtubule
KIF7
SUFU
GLI
Jagged
Delta
Notch
Notch signaling pathway
Fumarate
FH
Malate
Mitochondrion
HPH
HIF-1 signaling pathway
HIF-α
HIF-β
p300/CBP
Elongin B
Elongin C
CUL2
VHL
Rbx1
Oxidative or electrophilic stress
Acro
O2-
HO2
4HNE
H2O2
OH
NO
KEAP1
NRF2
CASP3/7
CASP9
Apaf-1
Cyt C
Apoptosis
Mitochondrion
FasL
Fas
FADD
CASP8
Bid
Bak
Bcl-2
Bcl-XL
NOXA
PUMA
Bad
Bim
TGFβ
TGFβRI
TGFβRII
Smad2/3
Smad4
TGF-β signaling pathway
Carcinogens
GSTP1
CASP3
DCC
CASP9
APPL
Adherens junction
α-catenin
ECAD
β-catenin
Tissue invasion & metastasis
Wnt
Frizzled
LRP5/6
GBP
Dvl
GSK-3β
Axin
APC
Wnt signaling pathway
Thrombin
LPA
Ang-II
GPCR
α12/13
RhoGEF
Rho
ROCK
SDF1
αi
AC
PGE2
αs
cAMP
PKA
cAMP signaling pathway
βγ
ECM
ITGA
ITGB
FAK
Focal adhesion
ECM-receptor interaction
BCR-ABL
CRKL
CBL
CRK
PTEN
PIP3
PI3K
NKX3.1
PKB/Akt
PI3K-Akt signaling pathway
BK
ET1
αq/11
PLCβ
IP3
DAG
Ca2+
PKC
Cytokine-cytokine receptor interaction
Cytokine
Receptor
Jak
TGFα
EGFR
EGF
ERBB2
PDGF
PDGFR
IGF
IGFR
KITLG
c-KIT
FLT3LG
FLT3
HGF
MET
FGF
FGFR
Grb2
Sos
RasGEF
EML4ALK
RET/PTC
RASSF1A
Ras
Raf
MEK
ERK
MAPK signaling pathway
RalGDS
Ral
RalBP1
Cdc42/Rac
PLD1
PA
Rac/Rho
JNK
TRK
Calcium signaling pathway
PLCγ
Elk
CAM
CAMK
IKK
IκBα
NFκB
mTOR
S6K
mTOR signaling pathway
CASP9
Bad
Bcl-XL
FKHR
MDM2
p53
p27
p21
Jak-STAT signaling pathway
STAT5
STAT3
STAT1
STAT2/4
Pim1/2
Bcl-XL
VEGF
NORE1A
RASSF1
MST1
CyclinD1
c-Jun
c-Fos
c-Myc
Ets1
MSK1
VEGF
MMPs
IL-8
CDK4
DAPK
BMP
GLI1
HIP1
PTCH1,2
HER2
Hes
Hey
VEGFR3
Glut1
VEGF
TGFα
PDGFβ
TGFβ
PSA
TCF/LEF
Survivin
CyclinD1
COX-2
iNOS
cIAPs
TRAFs
Bcl-2
Sustained angiogenesis
VEGF signaling pathway
Evading apoptosis
Proliferation
HO-1
NQO1
GSTs
TrxR1
BRCA2
Rad51
DNA
hMLH1
hMSH2
EVI1
AML-EVI1
CtBP
HDAC
Bax
hMSH3
hMSH6
TGFβRII
Insensitivity to anti-growth signals
Genomic damage
DNA damage
Genomic instability
p53 signaling pathway
p53
MITF
Resistance to chemotherapy
MDM2
p14/ARF
GADD45
Bax
Bak
p48
POLK
TERT
hTR
DNA
Telomeric DNA repeats
Immortality
p16INK4a
p21
Cell cycle
Rb
CDK4/6
CyclinD
p15INK4b
Miz1
c-Myc
Max
E2F
CDK2
CyclinE
p27/Kip1
CKS1
SCF
Skp2
Block of differentiation
GMCSFR
GCSFR
MCSFR
GCSFR
E2F
IL-6
c-Myc
MCSFR
TCR
RARα target genes
PU.1 target genes
C/EBPα target genes
AML1 target genes
PU.1
C/EBPα
AML1
PLZFRARα
PMLRARα
AML1ETO
CyclinA1
Estrogen signaling pathway
CyclinD1
c-Myc
PPARδ
ER
CoR
AP1/SP1
E2
TCF
β-catenin
RARβ
RXR
Retinoic acid
PPARγ
PPFP
Ligands
PPAR signaling pathway
05200 4/11/18
(c) Kanehisa Laboratories

**Suppl. Figure 4**. **The top significantly enriched KEGG pathway 'Pathways in Cancer' in aged compared with young adult human skeletal muscle tissue**. This pathway includes well known pathways in skeletal muscle regulation such as: Focal adhesion, MAPK signaling, PI3K-Akt-mTOR signaling, p53 signalling, Jak-STAT signaling, TGF-beta and Notch signalling. In this pathway 266 out of 277 CpG's were hypermethylated (RED) and only the remaining 11 hypomethylated (GREEN) in aged compared young adult muscle tissue. Note, because some genes have multiple CpGs per gene, Partek Pathway (Partek Genomics Suite) used to create this figure, selects the CpG with the lowest p-value (most significant) and uses the beta-value for that particular CpG to colour the pathway diagram. This is accurate therefore in situations where CpG's on the same gene have the same methylation pattern (e.g. all hyper or hypomethylated). However, where multiple CpG's on the same gene have a different methylation pattern (e.g. some hypomethylated and others hypermethylated), only the most significant methylated CpG is chosen and represented in the image. However, in this instance the majority of CpG's (even when there was more than one CpG per gene) were hypermethylated, so in this case the figure depicts an accurate representation of the data. Full CpG lists including the sites that were hypo and hypermethylated in this pathway are included in Suppl. File 1h.

**Title:** DNA methylation across the genome in aged human skeletal muscle tissue and stem cells: The role of HOX genes and physical activity

**Authors:** Turner DC, Gorski PP, Maasar MF, Seaborne RA, Baumert P, Brown AD, Kitchen MO, Erskine RM, Dos-Remedios I, Voisin S, Eynon N, Sultanov RI, Borisov OV, Larin AK, Semenova EA, Popov DV, Generozov EV, Stewart CE, Drust B, Owens DJ, Ahmetov II, Sharples AP.
