## Supplementary material for "DNA methylation across the genome in aged human skeletal muscle tissue and stem cells: The role of HOX genes and physical activity": Suppl. Figure 6

6a

‘Cytoskeletal protein binding’
0 h aged vs. 0 h young adult cells

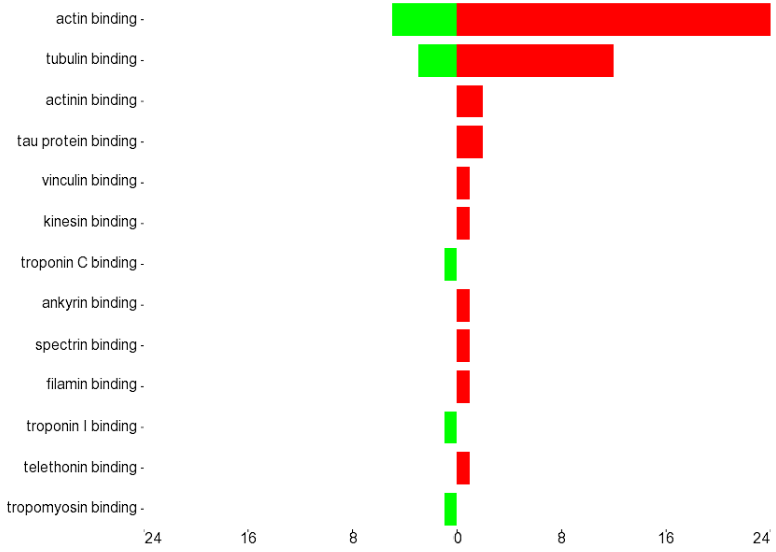


6b

‘Axon guidance’
0 h aged vs. 0 h young adult cells

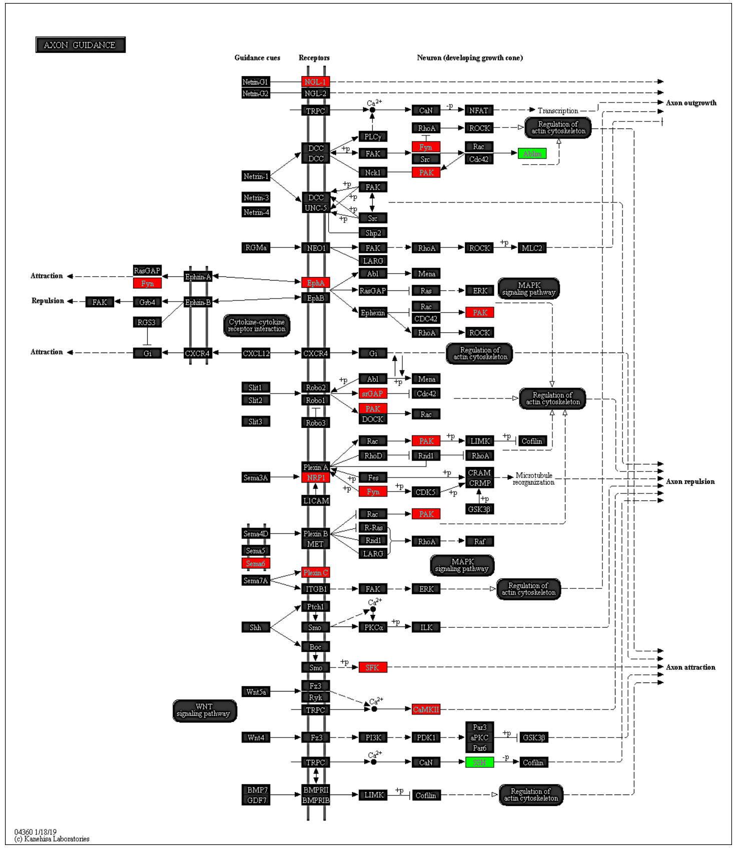


6c

‘Cytoskeletal protein binding’
72 h aged vs. 72 h young adult cells

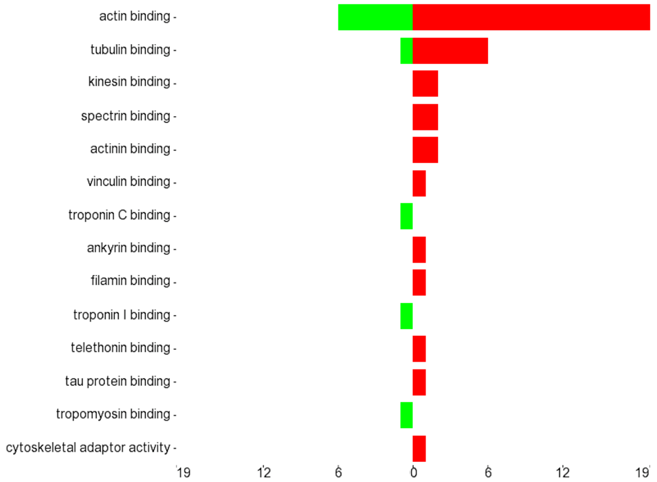


6d

‘cAMP signaling’
72 h aged vs. 72 h young adult cells

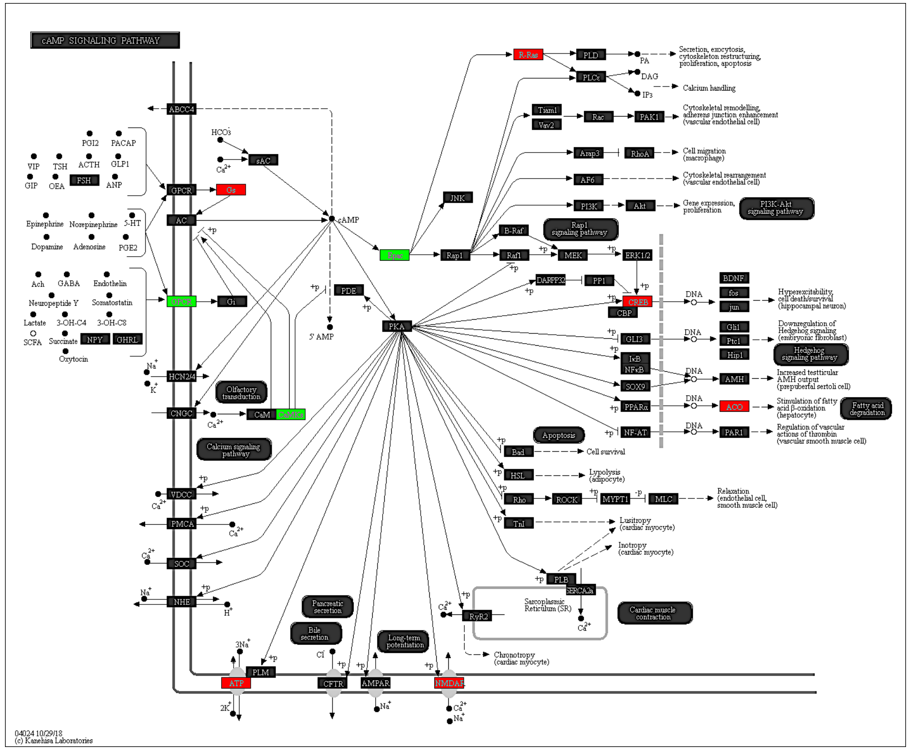

**Suppl. Figure 6. a.** 0 h aged vs. 0 h young adult muscle stem cells, top significantly enriched GO term 'cytoskeletal protein binding'. **b.** 0 h aged vs. 0 h young adult, top KEGG pathway 'axon guidance'. **c.** 72 h aged vs. 72 h young adult top GO term, 'cytoskeletal protein binding', **d.** 72 h aged vs. 72 h young adult, a top 3 KEGG pathway 'cAMP signaling'. All demonstrating a predominance of hypermethylation in aged compared with young muscle stem cells for these GO terms and KEGG pathways. RED hypermethylated and GREEN hypomethylated CpG's. Note, because some genes have multiple CpGs per gene, Partek Pathway (Partek Genomics Suite) used to create this figure, selects the CpG with the lowest p-value (most significant) and uses the beta-value for that particular CpG to colour the pathway diagram. This is accurate therefore in situations where CpG's on the same gene have the same methylation pattern (e.g. all hyper or hypomethylated). However, where multiple CpG's on the same gene have a different methylation pattern (e.g. some hypomethylated and others hypermethylated), only the most significant methylated CpG is chosen and represented in the image. Full and accurate CpG lists including the sites that were hypo and hypermethylated in these pathways are included in Suppl. File **3n** (for Suppl. Figure 6b) and Suppl. File **4k** (for Suppl. Figure 6d).

**Title:** DNA methylation across the genome in aged human skeletal muscle tissue and stem cells: The role of HOX genes and physical activity

**Authors:** Turner DC, Gorski PP, Maasar MF, Seaborne RA, Baumert P, Brown AD, Kitchen MO, Erskine RM, Dos-Remedios I, Voisin S, Eynon N, Sultanov RI, Borisov OV, Larin AK, Semenova EA, Popov DV, Generozov EV, Stewart CE, Drust B, Owens DJ, Ahmetov II, Sharples AP.
