## Supplementary material for "DNA methylation across the genome in aged human skeletal muscle tissue and stem cells: The role of HOX genes and physical activity": Suppl. Figure 7

7a ‘Developmental process’ (incl. anatomical structure development’) 7 d aged vs. 7 d young adult cells

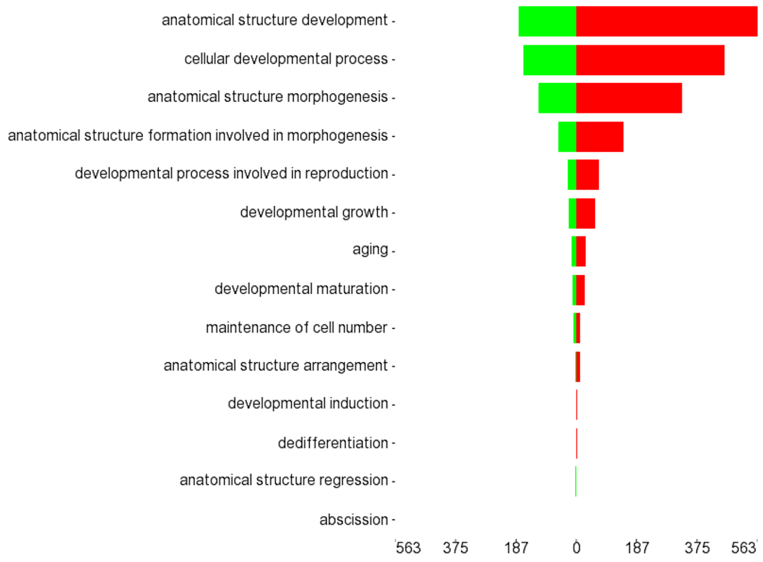


7b ‘PI3K-Akt Pathway’ 7 d aged vs. 7 d young adult cells

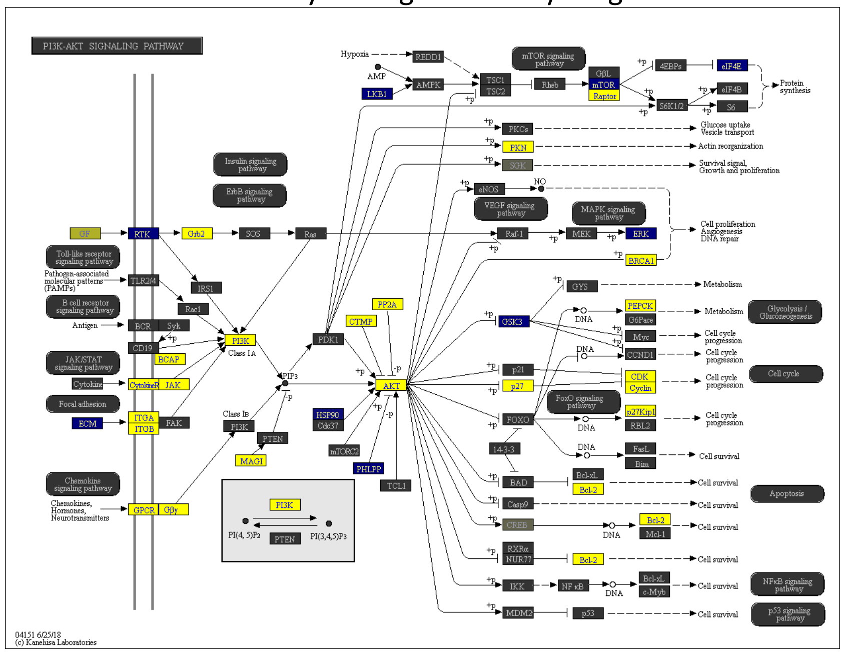


7c ‘GDP binding’ 10 d aged vs. 10 d young adult cells

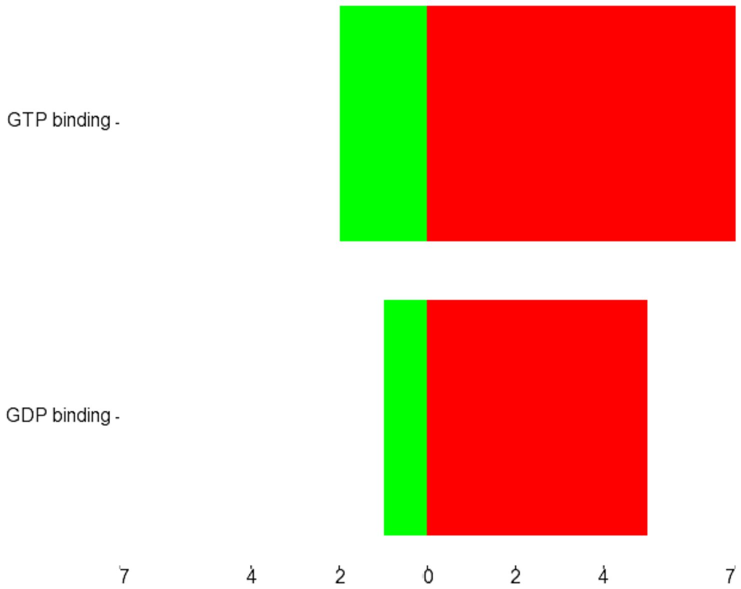


7d ‘Regulation of the actin cytoskeleton’ 10 d aged vs. 10 d young adult cells

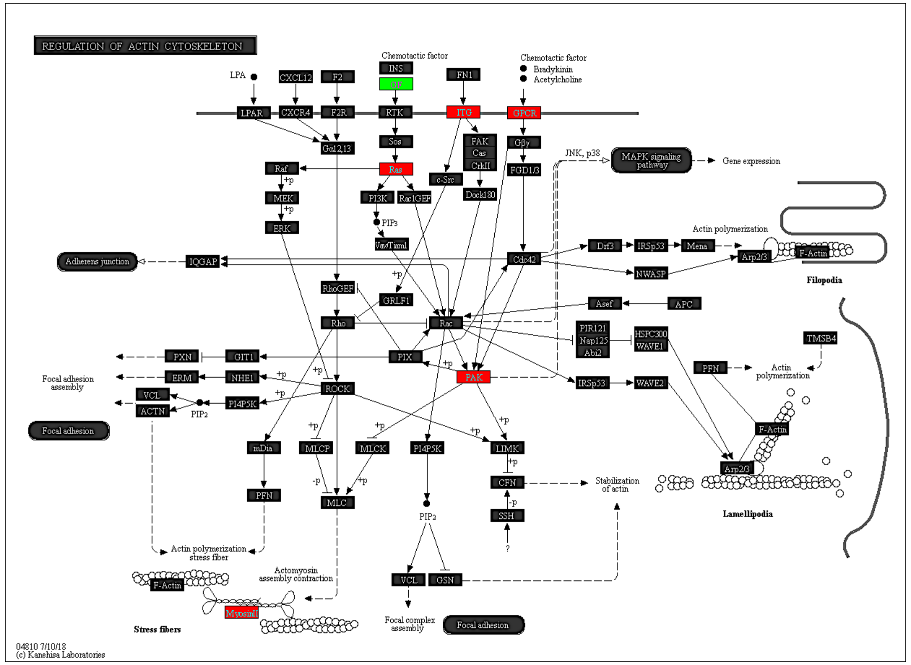

**Suppl. Figure 7. Comparison of DNA methylation in aged versus young muscle stem cells at 7 d and 10 d of differentiation. a.** 7 d aged vs. 7 d young adult skeletal muscle stem cells, top significantly enriched GO term, 'anatomical structure morphogenesis'. **b**. 7 d aged vs. 7 d young adult, top 10 KEGG pathway, 'PI3K/AKT pathway'..YELLOW hypermethylated, BLUE hypomethylated. **c**. 10 d aged vs 10 d young adult, top GO term, 'GDP binding'. **d.** 10 d aged vs. 10 d young adult, top KEGG pathway, 'regulation of the actin cytoskeleton'. RED hypermethylated and GREEN hypomethylated. Note, because some genes have multiple CpGs per gene, Partek Pathway (Partek Genomics Suite) used to create these pathway images, selects the CpG with the lowest p-value (most significant) and uses the beta-value for that particular CpG to colour the pathway diagram. This is accurate therefore in situations where CpG's on the same gene have the same methylation pattern (e.g. all hyper or hypomethylated). However, where multiple CpG's on the same gene have a different methylation pattern (e.g. some hypomethylated and others hypermethylated), only the most significant methylated CpG is chosen and represented in the image. Full and accurate CpG lists including the sites that are hypo and hypermethylated in these pathways are included in Suppl. File **5n** (for Suppl. Figure 7b) and Suppl. File **6i** (for Suppl. Figure 7d).

**Title:** DNA methylation across the genome in aged human skeletal muscle tissue and stem cells: The role of HOX genes and physical activity

**Authors:** Turner DC, Gorski PP, Maasar MF, Seaborne RA, Baumert P, Brown AD, Kitchen MO, Erskine RM, Dos-Remedios I, Voisin S, Eynon N, Sultanov RI, Borisov OV, Larin AK, Semenova EA, Popov DV, Generozov EV, Stewart CE, Drust B, Owens DJ, Ahmetov II, Sharples AP.
