## Supplementary material for "DNA methylation across the genome in aged human skeletal muscle tissue and stem cells: The role of HOX genes and physical activity": Suppl. Figure 8

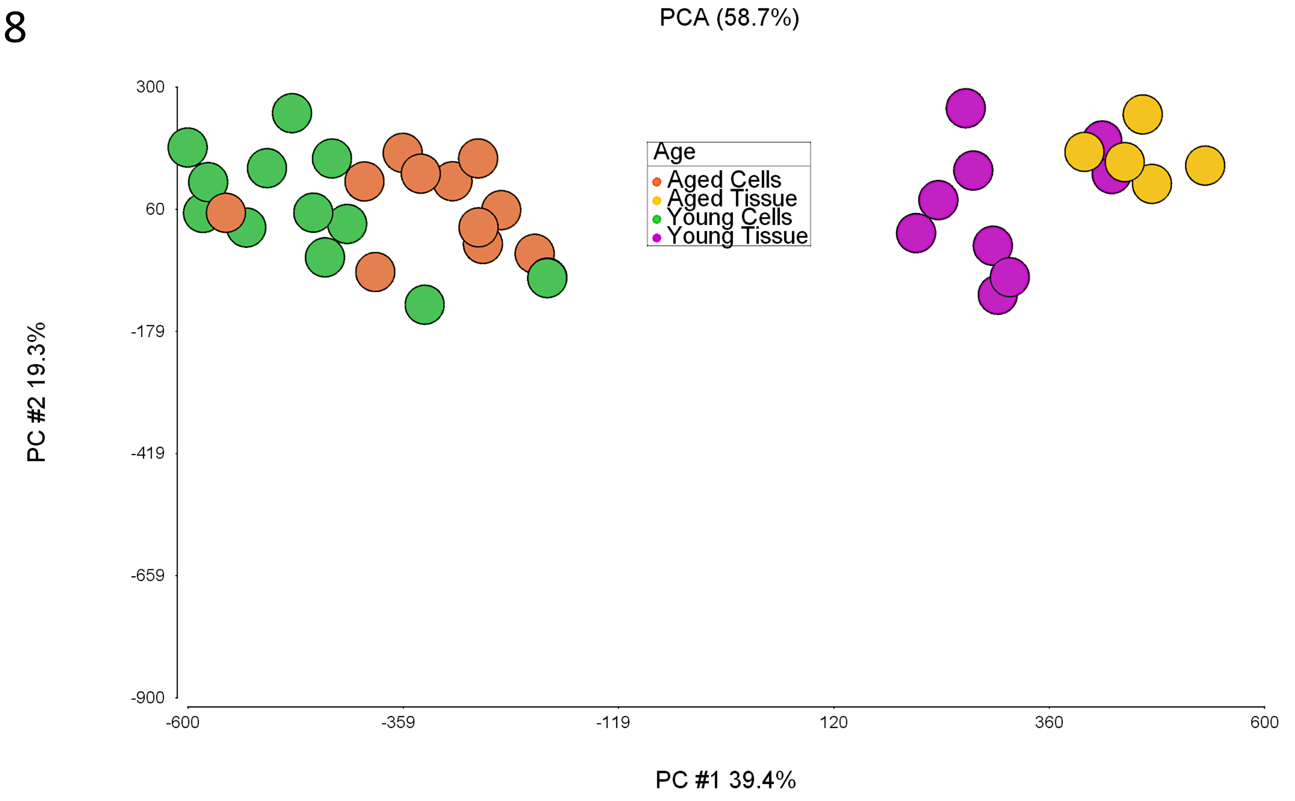


**Suppl. Figure 8.** PCA of aged and young adult skeletal muscle tissue versus aged and young adult isolated muscle stem cells across 0, 72 hr, 7 and 10 d of differentiation. Demonstrating methylation profiles of isolated muscle stem cells, even at late stages of differentiation, are vastly different compared with skeletal muscle tissue methylation profiles.

**Title:** DNA methylation across the genome in aged human skeletal muscle tissue and stem cells: The role of HOX genes and physical activity

**Authors:** Turner DC, Gorski PP, Maasar MF, Seaborne RA, Baumert P, Brown AD, Kitchen MO, Erskine RM, Dos-Remedios I, Voisin S, Eynon N, Sultanov RI, Borisov OV, Larin AK, Semenova EA, Popov DV, Generozov EV, Stewart CE, Drust B, Owens DJ, Ahmetov II, Sharples AP.
