## Supplementary material for "DNA methylation across the genome in aged human skeletal muscle tissue and stem cells: The role of HOX genes and physical activity": Suppl. Figure 10

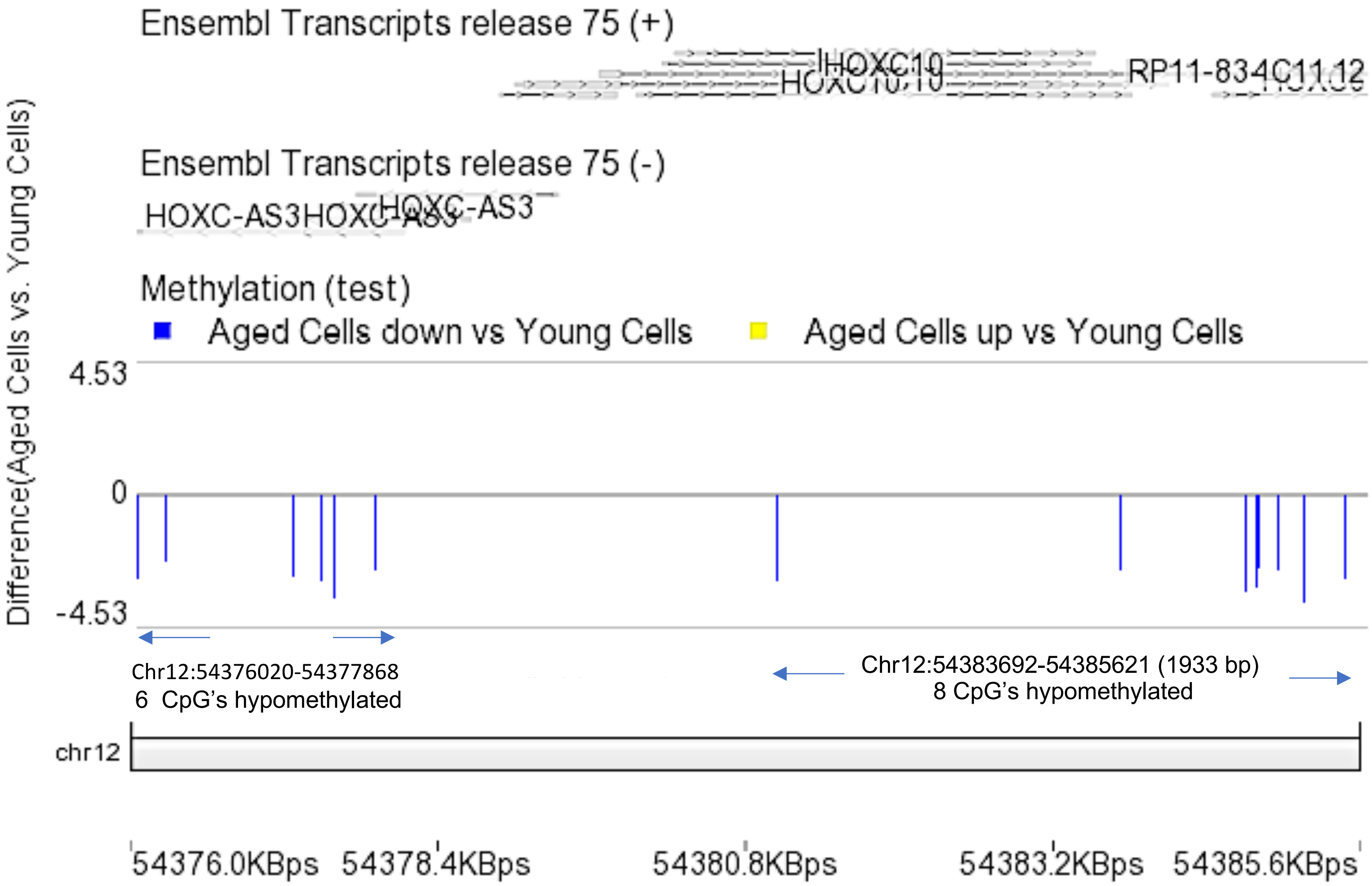


**Suppl. Figure 10. Differentially methylated region (DMR) analysis using** using the Bioconductor package DMRcate (DOI: 10.18129/B9.bioc.DMRcate) via Partek Genomics Suite. Hypomethylation of the HOXC10 (Chr12:54383692-54385621, 1933 bp) occurred in 8 CpG's and on the same chromosome just upstream of the HOXC10 gene (Chr12:54376020-54377868, 1849 bp) where there were another 6 CpG's hypomethylated within the lncRNA HOXC-AS3, in aged compared with young adult muscle stem cells. We also demonstrated that both HOXC10 (on average, not significantly) and HOXC-AS3 (significantly) increased in gene expression at 7 days of differentiation in aged compared with young adult muscle stem cells.

**Title:** DNA methylation across the genome in aged human skeletal muscle tissue and stem cells: The role of HOX genes and physical activity

**Authors:** Turner DC, Gorski PP, Maasar MF, Seaborne RA, Baumert P, Brown AD, Kitchen MO, Erskine RM, Dos-Remedios I, Voisin S, Eynon N, Sultanov RI, Borisov OV, Larin AK, Semenova EA, Popo
DV, Generozov EV, Stewart CE, Drust B, Owens DJ, Ahmetov II, Sharples AP.
